# TET knockout cells transit between pluripotent states and exhibit precocious germline entry

**DOI:** 10.1101/2024.12.02.626356

**Authors:** Raphaël Pantier, Elisa Barbieri, Sara Gonzalez Brito, Ella Thomson, Tülin Tatar, Douglas Colby, Man Zhang, Ian Chambers

**Author notes:** Joint 1^st^ authors (Raphaël Pantier, Elisa Barbieri, Sara Gonzalez Brito).

## Abstract

TET1, TET2 and TET3 are DNA demethylases with critical roles in development and differentiation. To assess the contributions of TET proteins to cell function during early development, single and compound knockouts of *Tet* genes in mouse pluripotent embryonic stem cells (ESCs) were generated. Here we show that TET proteins are not required to transit between naïve, formative and primed pluripotency. Moreover, ESCs with double-knockouts of *Tet1* and *Tet2* or triple-knockouts of *Tet1, Tet2* and *Tet3* are phenotypically indistinguishable. These TET-deficient ESCs exhibit differentiate defects; they fail to activate somatic gene expression and retain expression of pluripotency transcription factors. Therefore, TET1 and TET2, but not TET3 act redundantly to facilitate somatic differentiation. Importantly however, TET-deficient ESCs can differentiate into primordial germ cell-like cells (PGCLCs), and do so at high efficiency in the presence or absence of PGC-promoting cytokines. Moreover, acquisition of a PGCLC transcriptional programme occurs more rapidly in TET-deficient cells. These results establish that TET proteins act at the juncture between somatic and germline fates: without TET proteins, epiblast cell differentiation defaults to the germline.

## Introduction

Ten eleven translocation (TET) proteins promote active DNA demethylation by catalysing the oxidation of 5-methylcytosine (Tahiliani *et al*, 2009; Ito *et al*, 2011; He *et al*, 2011). In mammals, TET proteins are encoded by three genes (*Tet1*, *Tet2* and *Tet3)* which share an evolutionary conserved catalytic domain. TET proteins interact with multiple transcription factors (Costa *et al*, 2013; Okashita *et al*, 2014; Rampal *et al*, 2014; Wang *et al*, 2015; Sardina *et al*, 2018; Pantier *et al*, 2020), which could mediate their targeting to enhancer elements (Lu *et al*, 2014; Hon *et al*, 2014; Bogdanović *et al*, 2016; Ginno *et al*, 2020; Charlton *et al*, 2020). Additionally, TETs interact with proteins modulating chromatin, such as O-glycosyltransferase (Vella *et al*, 2013; Deplus *et al*, 2013; Shi *et al*, 2013; Chen *et al*, 2013) and the SIN3A co-repressor complex (Williams *et al*, 2011; Chandru *et al*, 2018; Zhu *et al*, 2018). Such partner protein interactions may be important in mediating the critical role of TET proteins during embryonic development.

TET proteins are expressed during early embryonic development and in ESCs (Ito *et al*, 2010; Gu *et al*, 2011; Dawlaty *et al*, 2011; Koh *et al*, 2011). In mice deletion of individual *Tet* genes is compatible with normal embryonic development but does lead to tissue specific phenotypes during later foetal development and in adults (Dawlaty *et al*, 2011; Li *et al*, 2011; Gu *et al*, 2011; Yamaguchi *et al*, 2012). In particular, TET1 deficiency leads to defects during later germline development, including incomplete reprogramming of genomic imprints and defective oocyte meiosis (Yamaguchi *et al*, 2012, 2013; Hackett *et al*, 2013; SanMiguel *et al*, 2018; Hill *et al*, 2018). Loss of TET2 causes defects in hematopoietic stem cell differentiation, leading predominantly to myeloid malignancies in adults (Li *et al*, 2011; Quivoron *et al*, 2011; Moran-Crusio *et al*, 2011; Ko *et al*, 2011; Muto *et al*, 2014). TET3 loss causes neonatal lethality, potentially due to its function in the reprogramming of the paternal genome in fertilized zygotes (Gu *et al*, 2011; Wang *et al*, 2013; Tsukada *et al*, 2015). In addition, the combined loss of TET1 and either TET2 or TET3 can cause morphological abnormalities and growth defects from embryonic day E10.5 (Dawlaty *et al*, 2013; Kang *et al*, 2015).

In contrast, embryos with triple knockouts of *Tet1, Tet2* and *Tet3* die shortly after implantation with gastrulation defects (Dai *et al*, 2016; Li *et al*, 2016). Consistent with this *in vivo* phenotype, ESCs with combined deletions of *Tet1, Tet2* and *Tet3* are blocked in somatic differentiation, whereas ESCs with single knockouts of *Tet* genes differentiate normally (Dawlaty *et al*, 2014; Verma *et al*, 2018). These studies indicate functional redundancy amongst TET proteins during early development of somatic cells. However, the abilities of cells lacking TET proteins to differentiate to the germline has not been assessed. Here, we removed all *Tet* family genes, singly and in combination, and performed *in vitro* differentiation assays to monitor the relative contributions of TET1, TET2 and TET3 to cellular transitions during peri-implantation development. Comparative analyses of cells with single, double and triple *Tet* gene deletions confirmed the somatic differentiation block in TET-deficient cells. Our results extend previous findings by showing that the somatic differentiation block is not due to an inability of TET-deficient cells to transit from naïve, through formative to primed pluripotency. Our results also establish that TET deficient cells are able to differentiate into the germline at high efficiency and do so at an accelerated rate and without the usual requirement for cytokine addition.

## Results

### Development of a novel strategy to knockout *Tet1/2/3* alleles by CRISPR/Cas9

Previous studies used a variety of approaches to knockout *Tet* alleles, including coding exon disruption by classic gene targeting (Li *et al*, 2011; Dawlaty *et al*, 2011; Gu *et al*, 2011; Yamaguchi *et al*, 2012; Zhang *et al*, 2013; Hu *et al*, 2014; Dai *et al*, 2016) or frameshift mutagenesis using CRISPR/Cas9 (Wang *et al*, 2013; Lu *et al*, 2014; Verma *et al*, 2018; Ginno *et al*, 2020). However, these strategies left open the formal possibility of truncated TET protein fragment expression. Therefore, to eliminate all endogenous *Tet1*, *Tet2* and *Tet3* coding potential the *Tet* open reading frames were excised using CRISPR/Cas9 and two sgRNAs targeting the start and stop codons, respectively (Figure EV1A-F). Single and combined *Tet* knockout cell lines were generated from wild-type E14Tg2a ESCs using a sequential strategy (Figure EV1G). Since our previous analyses indicated that TET3 protein is undetectable in ESCs (Pantier *et al*, 2019), we focussed our analysis of compound mutants on a comparison of *Tet1^-/-^*, *Tet2^-/-^* double knockout (DKO) and *Tet1^-/-^*, *Tet2^-/-^, Tet3^-/-^* triple knockout (TKO) cells. Targeted ESC lines were first assessed for expression of *Tet* mRNAs (Figure EV1H). As expected, expression of *Tet1, Tet2 and Tet3* mRNAs was lost in ESCs carrying deletions of *Tet1*, *Tet2* and *Tet3*, respectively.

### TET1 and TET2, but not TET3, are required for somatic lineage commitment

To assess the ability of *Tet* knockout ESCs to undergo somatic differentiation two distinct protocols were used: monolayer neural differentiation and embryoid body formation. Following monolayer neural differentiation (Ying *et al*, 2003), flattened differentiated cells appeared in wild-type, *Tet2^-/-^* and *Tet3^-/-^* cultures. In contrast, colonies with compact, rounded morphologies resembling ESC colonies persisted in *Tet1*^-/-^ single knockout*, Tet1^-/-^*, *Tet2^-/-^* DKO and *Tet1^-/-^*, *Tet2^-/-^, Tet3^-/-^* TKO cultures (Figure EV2A). After 8 days of differentiation, *Tet1*^-/-^ single knockout*, Tet1^-/-^*, *Tet2^-/-^* DKO and *Tet1^-/-^*, *Tet2^-/-^, Tet3^-/-^* TKO cultures also retained mRNAs encoding the pluripotency transcripts *Oct4*, *Esrrb* and *Rex1* (Figure EV2B). Compared to wild-type, *Tet2^-/-^*and *Tet3^-/-^* cell lines, *Tet1^-/-^* cells showed reduced induction of the neural markers *Sox1*, *Ascl1*, *Tuj1* and *N-cadherin* (Figure EV2C). *Ascl1*, *Tuj1* and *N-cadherin* mRNAs were reduced further in *Tet1^-/-^*, *Tet2^-/-^* DKO and *Tet1^-/-^*, *Tet2^-/-^, Tet3^-/-^* TKO cells (Figure EV2C). Consistent with these changes TUJ1-positive neurons were undetectable in *Tet1*^-/-^ single knockout, *Tet1^-/-^*, *Tet2^-/-^* DKO and *Tet1^-/-^*, *Tet2^-/-^, Tet3^-/-^* TKO cultures (Figure EV2D). These results show that the functions of TET1 and, to a lesser extent, TET2 are required in combination for efficient neural differentiation.

Multi-lineage differentiation was assessed after embryoid body (EB) formation. This showed the persistence of some colonies with an undifferentiated ESC-like morphology after 14 days of EB culture of *Tet1^-/-^*, *Tet2^-/-^* DKO and *Tet1^-/-^*, *Tet2^-/-^, Tet3^-/-^* TKO cells (Figure EV3A). In contrast, wild-type or single knockout *Tet* cultures appeared fully differentiated (Figure EV3A). In addition, after 14 days *Tet1^-/-^*, *Tet2^-/-^*DKO and *Tet1^-/-^*, *Tet2^-/-^, Tet3^-/-^*TKO EBs retained expression of the pluripotency transcription factors *Oct4*, *Esrrb* and *Nanog* (Figure EV3B) and failed to upregulate the germ layer markers *Gata4*, *Col1a1, Flk1* and *Sox17* (Figure EV3C). Furthermore, *Tet1^-/-^*, *Tet2^-/-^* DKO and *Tet1^-/-^*, *Tet2^-/-^, Tet3^-/-^* TKO EBs also showed an almost complete lack of beating colonies, indicating a block to mature cardiomyocyte differentiation (Figure EV3D).

Together, these results extend previous analysis by showing that it is the combined actions of TET1 and TET2, and not TET3, that are required for multi-lineage specification.

### The combined loss of TET1 and TET2 enhances ESC self-renewal

To examine the effect of *Tet* gene deletions on the undifferentiated ESC phenotype, we assessed the morphology, self-renewal efficiency and expression of pluripotency markers of *Tet* knockout ESC lines. At routine passaging density, phase contrast imaging of live ESC cultures indicated that loss of TET proteins did not affect ESC morphology (Figure EV4A). To quantitatively assess self-renewal efficiency, *Tet* knockout ESCs were plated at clonal density, cultured for 7 days in the presence or absence of leukemia inhibitory factor (LIF) and stained for alkaline phosphatase activity (Figure 1A and EV4B). Interestingly, both *Tet1^-/-^*, *Tet2^-/-^* DKO and *Tet1^-/-^*, *Tet2^-/-^, Tet3^-/-^* TKO ESCs showed a strong increase in the proportion of undifferentiated colonies formed in the presence of LIF (Figure 1A). However, no differences were seen in the colonies formed by any of these lines in the absence of LIF (Figure EV4B). This increased responsiveness to LIF was reflected in the morphology of individual colonies formed by *Tet1^-/-^*, *Tet2^-/-^* DKO and *Tet1^-/-^*, *Tet2^-/-^, Tet3^-/-^* TKO ESC clones, which were more undifferentiated than wild-type or single *Tet* knockout ESCs, with fewer differentiated cells present at the edges (Figure 1B). Together these results indicate that the redundant activities of TET1 and TET2 reduce the efficiency of ESC self-renewal.

**Figure 1.**
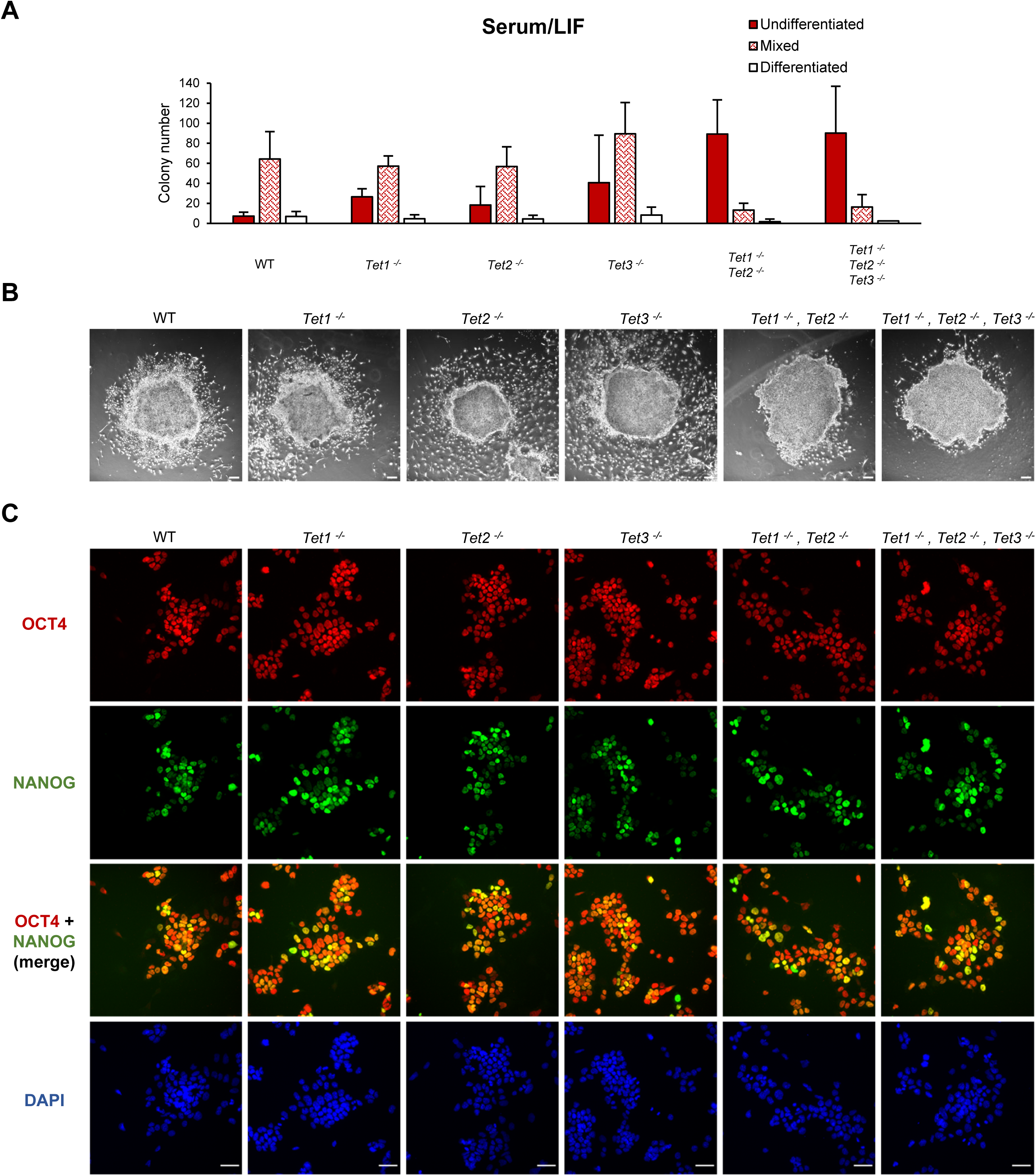
Characterisation of TET function during ESC self-renewal. **A.** Alkaline phosphatase staining of the indicated ESC lines following plating at clonal density in serum/LIF medium for 7 days. Colonies were counted and categorised according to their alkaline phosphatase staining. Error bars: standard deviation in 4 independent replicate experiments. **B.** For each of the genotypes quantified in (A) an image representative of the main class of colonies formed in serum/LIF is shown by phase contrast microscopy. Scale bars: 100µm. **C.** Co-immunofluorescence for OCT4 (red) and NANOG (green) in the indicated ESC lines cultured in serum/LIF. Scale bars: 50µm.

However, co-immunofluorescence analysis indicated that single and combined *Tet* gene deletions did not affect the levels and distribution of the pluripotency factors NANOG and OCT4 in cell cultures (Figure 1C). Notably, NANOG remains heterogeneously expressed in *Tet* mutant lines, suggesting that loss of TET proteins may not affect transitions between the pluripotent cell states present in serum-containing cultures (Chambers *et al*, 2007).

### Loss of TETs does not affect transitions from naïve to formative or primed pluripotency

To further assess the role of TET proteins during the transitions between pluripotent states, *Tet* knockout ESCs were differentiated into epiblast-like cells (EpiLCs) (Hayashi *et al*, 2011; Hayashi & Saitou, 2013). All *Tet* mutant cell lines differentiated into EpiLCs within 48h, without noticeable morphological differences between wild-type and *Tet* knockout lines (Figure EV4C). To assess transcriptional changes accompanying the transition from naïve to formative pluripotency, RNA was analysed at 0h and 48h of EpiLC differentiation. In *Tet* knockout and wild-type cells, the levels of the naïve pluripotency mRNAs *Esrrb*, *Prdm14* and *Rex1* rapidly decreased, while *Oct4* mRNA remained expressed (Figure 2A). Conversely, the formative pluripotency markers *Otx2* and *Oct6* were upregulated in a similar manner in wild-type and *Tet* knockout EpiLCs (Figure 2B). These results indicate that TET proteins are dispensable for the transition from naïve to formative pluripotency.

**Figure 2.**
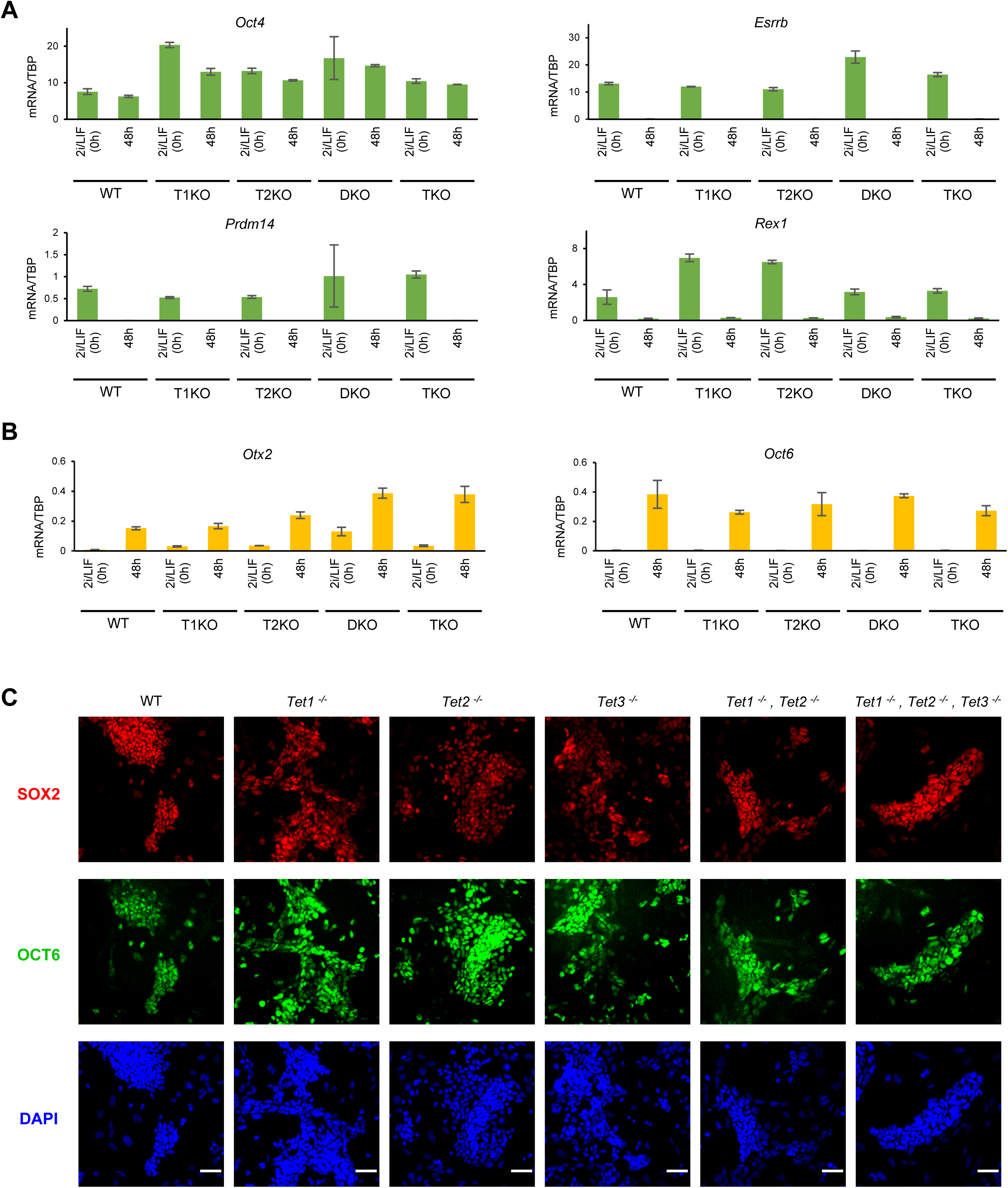
Characterisation of TET function during the transition from naïve to primed pluripotency. **A, B.** Pluripotency mRNA levels at 0 and 48 hours of EpiLC differentiation in the indicated cell lines. mRNA levels were quantified by RT-qPCR and normalised to TBP mRNA levels. Error bars: standard deviation in 3 technical replicates. **C.** Co-immunofluorescence for SOX2 (red) and OCT6 (green) in the indicated EpiSC lines (passage 11) cultured in activin/FGF. Scale bars: 50µm.

To assess the requirement for TET protein function for the transition to primed pluripotency, *Tet* knockout ESCs were differentiated into epiblast stem cells (EpiSCs) *in vitro* (Guo *et al*, 2009). Stable self-renewing EpiSC lines lacking either single or combined *Tet* alleles were obtained. Phase contrast microscopy indicated that all mutant lines had a typical flat, elongated morphology (Figure EV4D). Co-immunofluorescence analysis confirmed that the primed pluripotency factors OCT6 and SOX2 were expressed in all *Tet* knockout EpiSC lines, similarly to wild-type cells (Figure 2C). As EpiSCs resemble epiblast cells in early post-implantation embryos at ∼E5.5-6.5 (Tesar *et al*, 2007; Brons *et al*, 2007; Han *et al*, 2010), these results are compatible with the fact that *Tet1^-/-^*, *Tet2^-/-^, Tet3^-/-^* triple knockout embryos are undistinguishable from wild-type at this developmental stage (Dai *et al*, 2016; Li *et al*, 2016). Together, our results extend previous reports that *Tet1^-/-^*, *Tet2^-/-^, Tet3^-/-^* triple knockout cells are impaired in somatic differentiation *in vitro* (Dawlaty *et al*, 2014; Li *et al*, 2016; Verma *et al*, 2018) by showing that this is also the case for *Tet1^-/-^*, *Tet2^-/-^* double knockout cells and by showing that *Tet* mutants are unimpaired in their ability to transit between pluripotent states.

### TET-deficient cells efficiently acquire germline surface markers without requiring inductive cytokines

To assess the ability of *Tet* mutant cells to enter germline development, wild-type and *Tet* knockout ESCs were transitioned to an EpiLC state before further differentiation into primordial germ cell-like cells (PGCLCs) (Hayashi *et al*, 2011; Hayashi & Saitou, 2013). After 6 days, wild-type and single *Tet* deleted lines produced similar low levels of cells displaying the PGCLC surface markers SSEA-1 and CD61 (Figure 3A, C). In contrast, the proportion of SSEA1^+^/CD61^+^ cells was strongly increased in both *Tet1^-/-^, Tet2^-/-^* DKO and *Tet1^-/-^, Tet2^-/-^, Tet3^-/-^* TKO lines (Figure 3A, C). This result is reminiscent of the increased proportion of SSEA1^+^/CD61^+^ cells formed in cells deleted for the transcription factor *Otx2* (Zhang *et al*, 2018a). In the case of *Otx2*-null cells, ∼30% of cells could even form SSEA1^+^/CD61^+^ cells without the usual requirement for PGC-promoting cytokines. To determine if this was also the case for *Tet* mutants, differentiation was performed in the absence of cytokines. Surprisingly, both *Tet1^-/-^, Tet2^-/-^* DKO and *Tet1^-/-^, Tet2^-/-^, Tet3^-/-^*TKO lines yielded SSEA1^+^/CD61^+^ cells at similar efficiencies regardless of the presence or absence of cytokines, thereby outperforming *Otx2*-null cells in this regard (Figure 3B,C). These results suggest that the combined loss of TET1 and TET2 proteins enables enhanced PGCLC differentiation and importantly permits this differentiation to proceed without the requirement for cytokine signalling.

**Figure 3.**
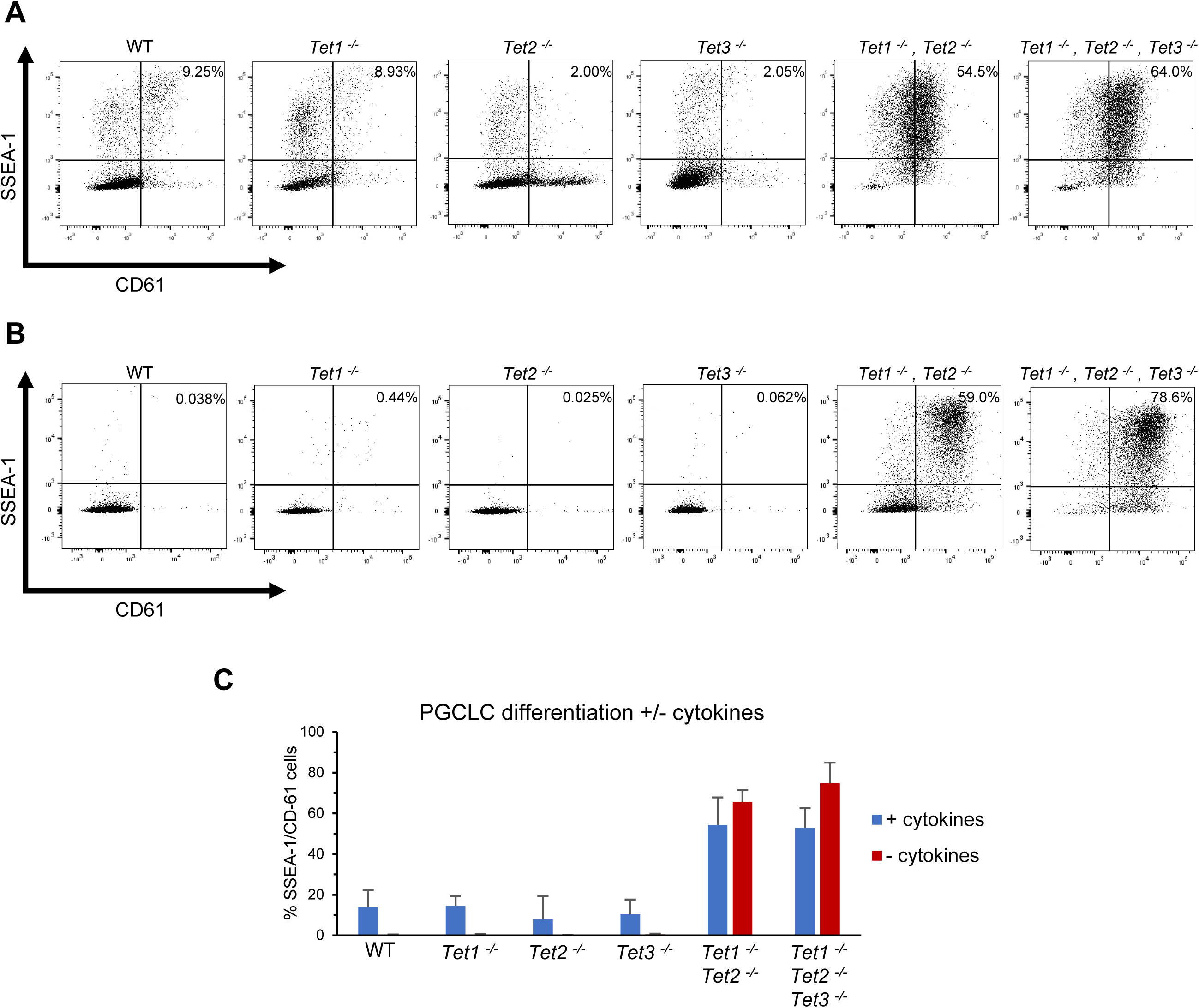
Characterisation of TET function during germline commitment. **A, B.** Flow cytometric analysis of the indicated cell lines following 6 days of PGCLC differentiation with (A) or without (B) PGC-promoting cytokines. SSEA-1/CD61 double-positive cells on the top right quadrant report PGCLCs. **C.** Quantification of SSEA-1/CD61 double-positive cells following 6 days of PGCLC differentiation with (blue) or without (red) cytokines in the indicated cell lines. Error bars: standard deviation in 3 independent replicate experiments.

### Transcriptional changes indicate efficient germline differentiation of TET-deficient cells

To determine whether the appearance of the PGCLC markers CD61 and SSEA1 on the surface of TET-deficient cells was reflected in transcriptional changes consistent with PGC differentiation, RNA-seq analysis was performed on wild-type and *Tet1^-/-^, Tet2^-/-^, Tet3^-/-^* TKO cells in both EpiLCs and during PGCLC differentiation. PGCLC samples were compared to a cell population known not to contain PGCLCs, obtained by differentiating wild-type cells without PGC-promoting cytokines. First, *Tet* gene expression dynamics during PGCLC differentiation was analysed. In wild-type cells *Tet1* and *Tet2* are upregulated in cultures undergoing germline differentiation in the presence of cytokines but are lowly expressed in cultures in the absence of cytokines where no PGCLCs form. In contrast, *Tet3* is downregulated in the presence of cytokines but upregulated in the absence of cytokines (Figure EV5A).

Principal component (PC) analysis of the RNA-seq data shows that wild-type and *Tet1^-/-^*, *Tet2^-/-^, Tet3^-/-^* TKO EpiLCs cluster together (Figure 4A). This global transcriptional analysis confirms the ability of *Tet1^-/-^*, *Tet2^-/-^, Tet3^-/-^* ESCs to transit to an EpiLC state (Figure 2; EV4C). From this common starting point, wild-type EpiLCs differentiated in the absence of cytokines move along both PCs (Figure 4A). In contrast, wild-type EpiLCs differentiated for 6 days in the presence of PGC-promoting cytokines show the opposite variation along PC1 (Figure 4A). Interestingly, after only 2 days *Tet1^-/-^*, *Tet2^-/-^, Tet3^-/-^* TKO PGCLCs are transcriptionally comparable to wild-type cells after 6 days of PGCLC differentiation (Figure 4A), and this is the case with or without PGC-promoting cytokines (Figure 4A). After 6 days, *Tet1^-/-^*, *Tet2^-/-^, Tet3^-/-^* TKO PGCLCs have diverged further along PC2, and this occurs in the presence or absence of cytokines. These results suggest that during PGCLC differentiation *Tet1^-/-^*, *Tet2^-/-^, Tet3^-/-^* TKO cells undergo similar changes to wild-type PGCLCs and that these changes do not require PGC-promoting cytokines.

**Figure 4.**
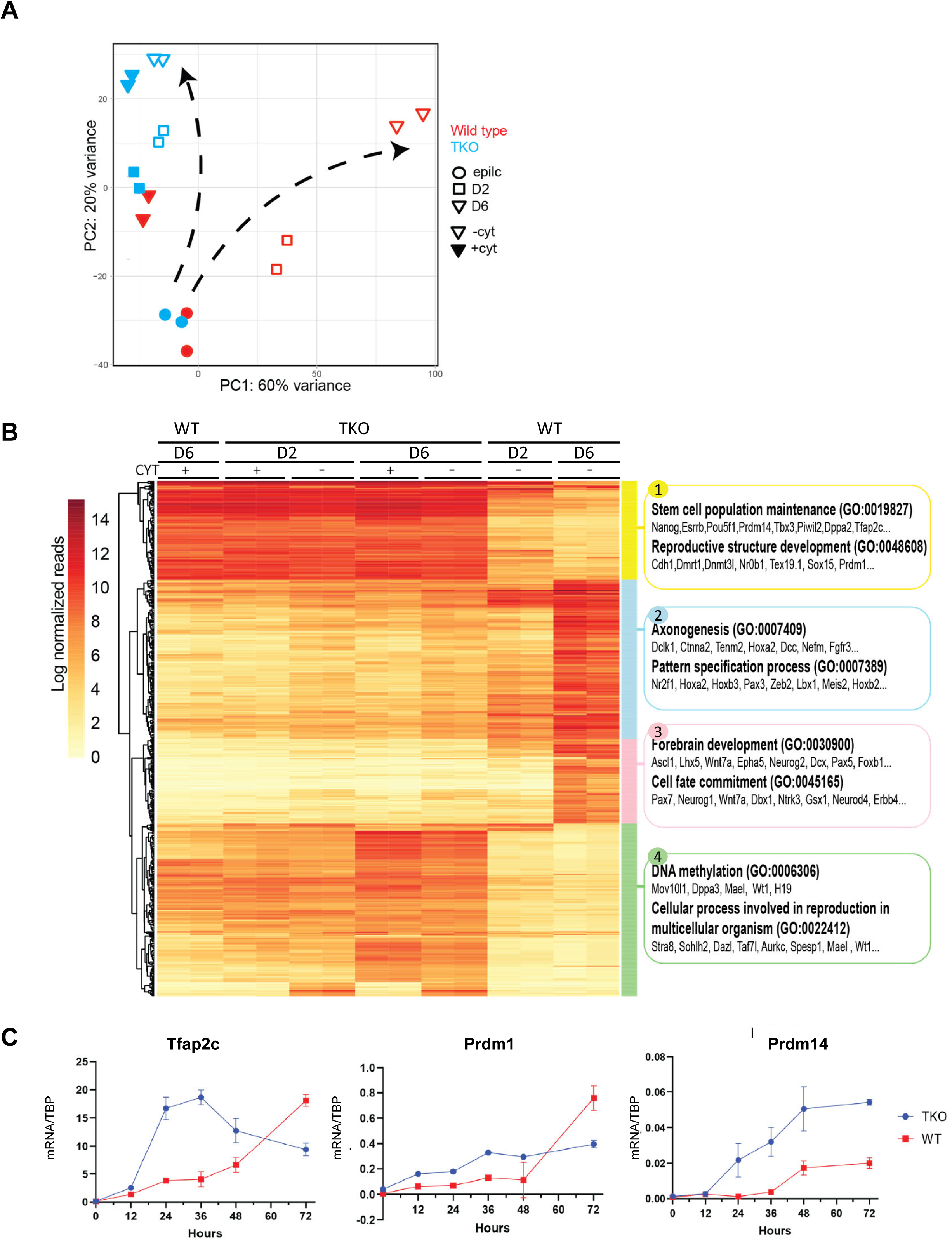
TET-TKOs upregulate germline markers in an accelerated manner. **A**. Principal component analysis (PCA) of bulk RNA-seq samples. Wild-type and TKO cells at day 0 (EpiLCs), day 2 and day 6 of differentiation (+/-cytokines). Wild-type cells differentiated in the presence of cytokines were sorted into SSEA1^+^/CD61^+^ populations to select for PGCLCs. All other populations were collected as full aggregates (n=2). **B**. Heatmap of the log normalized reads of the top variable genes across all samples (top 400 genes contributing to PC1 and top 400 genes contributing to PC2). Genes are grouped by hierarchical clustering and the top 2 GO terms are shown for each group (right hand side). **C**. Expression of key germline mRNAs measured at the indicated times of differentiation in the presence of cytokines. Error bars: standard deviation in 3 technical replicates

The above analysis suggests that by day 2, *Tet1^-/-^*, *Tet2^-/-^, Tet3^-/-^* TKO cells have a developmentally advanced transcriptome. To understand the global transcriptional changes behind the PCA the expression and biological function of the top 400 genes contributing to PC1 and PC2 was analysed (Figure 4A, B) and day 2 *Tet1^-/-^*, *Tet2^-/-^, Tet3^-/-^* TKO cells were directly compared to wild-type PGCLCs (Figure EV5 B - D).

The top 800 genes contributing to the PCA were grouped by hierarchical clustering resulting in four distinct groups (Figure 4B). Group 1 includes genes associated with “stem cell population maintenance” and “reproductive structure development”, such as *Nanog*, *Prdm14*, *Tfap2c* and *Prdm1.* Group 1 showed similar levels of expression throughout all PGCLCs (wild-type plus cytokines and *Tet1^-/-^*, *Tet2^-/-^, Tet3^-/-^*TKO plus and minus cytokines), but reduced levels of expression in wild-type cells differentiating without cytokines (Figure 4B). A comparison of the transcriptional signatures supports an overlap between day 6 wild-type cells and *Tet1^-/-^, Tet2^-/-^, Tet3^-/-^* TKO cells at day 2 (Figure EV5B, D). The majority of genes upregulated in wild-type PGCLCs at day 6 were also upregulated in *Tet1^-/-^, Tet2^-/-^, Tet3^-/-^* TKO PGCLCs as early as day 2, and this occurred irrespective of the presence or absence of cytokines (1080 genes, 76.5% of all WT genes) (Figure EV5B). Commonly upregulated genes are mostly related to reproduction and meiosis and include key germline regulators *Prdm14*, *Prdm1* and *Tfap2c,* which are expressed at similar levels in wild-type PGCLCs at day 6 and in *Tet1^-/-^, Tet2^-/-^, Tet3^-/-^* TKO PGCLCs at day 2 (Figure EV5C). Therefore, by day 2 *Tet1^-/-^, Tet2^-/-^, Tet3^-/-^* TKO PGCLCs have undergone the same upregulation of the germline program that takes 6 days to occur in wild-type PGCLCs.

Groups 2 and 3 show similar expression patterns. These genes are highly expressed in wild-type cells after 6 days of differentiation without cytokines but remain low in wild-type cells differentiated with cytokines and in all *Tet1^-/-^*, *Tet2^-/-^, Tet3^-/-^* cells analysed, irrespective of the presence or absence of cytokines (Figure 4B). Genes in groups 2 and 3 are associated with neural differentiation and include *Ascl1, Pax7, Neurog1* and *Neurog2* (Figure 4B). A comparative analysis of downregulated genes showed that most genes downregulated in wild-type PGCLC were already repressed in *Tet1^-/-^*, *Tet2^-/-^, Tet3^-/-^* PGCLCs by day 2, and again that this occurred irrespective of the presence or absence of cytokines (Figure EV5D). Commonly downregulated genes include neural markers like *Neurog2 and Neurod4* (Figure EV5C), and associated gene ontology terms are mostly related to the neural fate (Figure EV5D). This suggest that in the absence of cytokines wild-type cells follow a neural fate, which is repressed in *Tet1^-/-^*, *Tet2^-/-^, Tet3^-/-^* TKO cells regardless of the addition of cytokines.

Overall, these results suggest that by day 2 *Tet1^-/-^*, *Tet2^-/-^, Tet3^-/-^* TKO cells differentiated in the presence or absence of cytokines both upregulate the germline program and repress the somatic program.

### TET-deficient PGCLCs show precocious expression of late germline markers

A fourth group are expressed highly in *Tet1^-/-^*, *Tet2^-/-^, Tet3^-/-^* TKO PGCLCs after 6 days of differentiation (with or without cytokines) (Figure 4B). This group differs from group 1 by being expressed at lower levels in wild-type PGCLCs at day 6 than in *Tet1^-/-^*, *Tet2^-/-^, Tet3^-/-^* TKO PGCLCs. This reflects PC differences between wild-type and *Tet1^-/-^*, *Tet2^-/-^, Tet3^-/-^* TKO cells at day 6 (Figure 4A and 4B). Gene ontology (GO) enrichment shows that genes in group 4 are mostly associated with the gene ontology terms “DNA methylation” and includes *Mov10l1, Dppa3, Mael* and *Wt1*, genes previously associated with germline or gonad development (Sato *et al*, 2002; Wilhelm & Englert, 2002; Soper *et al*, 2008; Hackett *et al*, 2012; Vourekas *et al*, 2015) and “cellular process involved in reproduction”, and includes the late germline markers *Stra8, Dazl and Taf7l* (Figure 4B). Indeed, temporal assessment shows an increased expression of *Stra8*, *Dazl*, *Taf7l* and *Spesp1* mRNAs in *Tet1^-/-^*, *Tet2^-/-^, Tet3^-/-^* TKO PGCLCs as early as day 2 (Figure EV5E).

### Expression of PGC TFs indicates accelerated germline entry in TET-deficient cells

The expression of late markers suggests that the transcriptome of *Tet1^-/-^*, *Tet2^-/-^, Tet3^-/-^* TKO PGCLCs might reflect a later stage in PGC development, consistent with the accelerated germ cell differentiation pinpointed by the PC analysis. To assess whether the similarity between day 2 *Tet1^-/-^*, *Tet2^-/-^, Tet3^-/-^* TKO PGCLCs and day 6 wild type PGCLCs could be explained by accelerated germline entry, the expression of *Tfap2c, Prdm1* and *Prdm14* was examined at 12-hourly intervals during the first 3 days of PGCLC differentiation. Following the addition of cytokines, all three mRNAs show increased expression in *Tet1^-/-^*, *Tet2^-/-^, Tet3^-/-^* TKO cells over the first 2 days of differentiation, compared to a more delayed upregulation in wild-type cells (Figure 4C). In particular, *Tfap2c* and *Prdm14* mRNAs are more highly expressed in *Tet1^-/-^*, *Tet2^-/-^, Tet3^-/-^*TKO cells between 24 and 48 hours (Figure 4C).

Taken together, this data shows that germline markers are expressed at similar levels in *Tet1^-/-^*, *Tet2^-/-^, Tet3^-/-^*TKO PGCLCs and wild-type PGCLCs, but that somatic markers upregulated in wild-type cells in the absence of cytokines remain low in *Tet1^-/-^*, *Tet2^-/-^, Tet3^-/-^* TKO PGCLCs throughout differentiation. Moreover, *Tet1^-/-^*, *Tet2^-/-^, Tet3^-/-^* TKO PGCLCs enter the germline at an accelerated rate. The major differences between *Tet1^-/-^*, *Tet2^-/-^, Tet3^-/-^* TKO PGCLCs and wild-type PGCLCs at day 6 are in mRNAs characteristic of later germline development; differences that are not evident in day 2 *Tet1^-/-^*, *Tet2^-/-^, Tet3^-/-^* TKO PGCLCs. Alongside the earlier upregulation of *Prdm14, Tfap2c* and *Blimp1* mRNAs, this suggests precocious germline development of *Tet1^-/-^*, *Tet2^-/-^, Tet3^-/-^* TKO cells.

## Discussion

Here, we have compared ESC lines lacking one or several *Tet* alleles. This allowed us to characterise unique and redundant functions of TET proteins and to show the early cellular changes during distinct differentiation contexts. This uncovered key redundant roles for TET1 and TET2 in supporting somatic lineage differentiation and acting as a critical barrier to germline induction (Figure 5).

**Figure 5.**
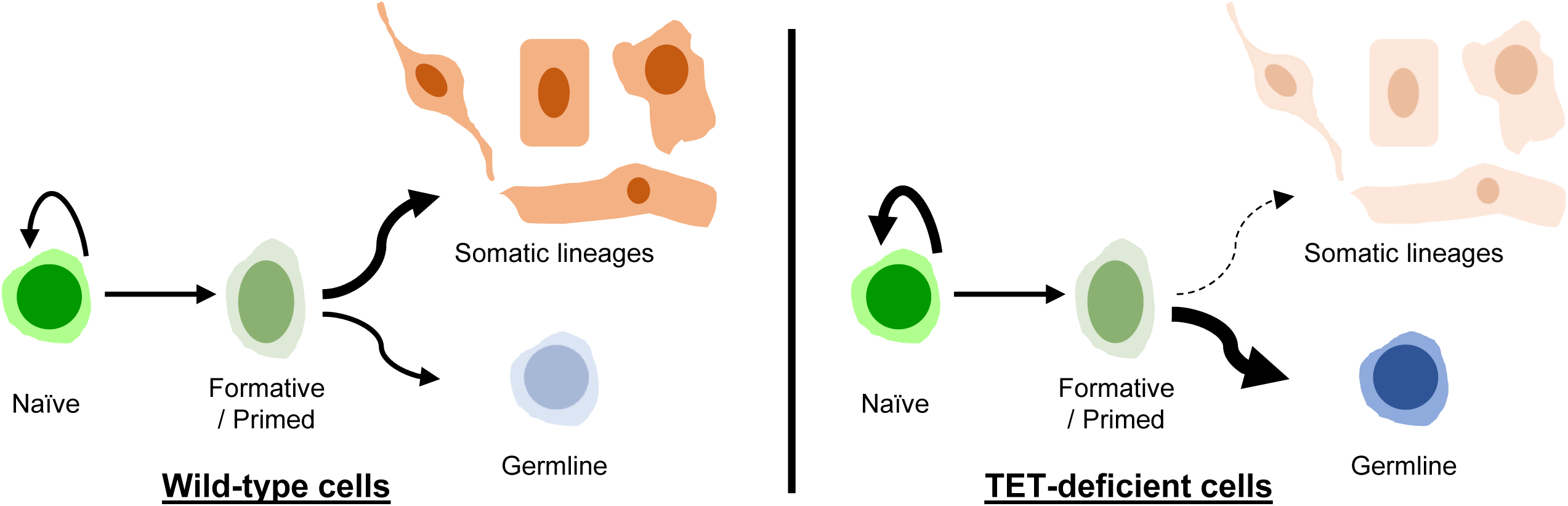
TET proteins control the choice between somatic and germline fates. Diagram summarising the phenotype observed in TET-deficient cells. *Tet1^-/-^, Tet2^-/-^* (DKO) and *Tet1^-/-^,Tet2^-/-^,Tet3^-/-^*(TKO) cells can transit between naïve, formative and primed pluripotency, in a similar way to wild type cells. However, these cells show an enhanced capacity to commit to a germline fate and fail to commit to somatic lineages.

During routine passaging in serum/LIF, all *Tet* knockout ESC lines have a normal morphology and express similar levels of pluripotency markers to wild-type ESCs. Furthermore, TET-deficient ESCs are defective in differentiation into somatic lineages and maintain expression of pluripotency genes. Both a block to differentiation and retention of pluripotency were previously observed in triple *Tet* knockout cells and were attributed to redundant activities of TET1, TET2 and TET3 (Dawlaty *et al*, 2014; Verma *et al*, 2018). However, we show here that *Tet1^-/-^, Tet2^-/-^* DKO and *Tet1^-/-^, Tet2^-/-^, Tet3^-/-^* TKO ESCs have similar blocks to differentiation. Therefore, redundancy amongst TET proteins in driving loss of pluripotency is restricted to TET1 and TET2 and does not extend to TET3.

*Tet1^-/-^, Tet2^-/-^* DKO and *Tet1^-/-^, Tet2^-/-^, Tet3^-/-^* TKO ESCs have more undifferentiated colony morphologies than wild-type or single *Tet* knockout cells when plated at clonal density in LIF. This appears to result from an enhanced sensitivity to LIF, rather than LIF independence, since undifferentiated colonies do not form when mutant cell lines are plated at clonal density in the absence of exogenous LIF. Whether this enhanced LIF sensitivity is connected to the recent observation that TET1 and TET2 are required for pluripotent cells to survive embryonic diapause (Stötzel *et al*, 2024), a process that also requires signalling through the LIFR/gp130 pathway (Nichols *et al*, 2001) will be an interesting area for future investigation. Notably, *Tet1^-/-^, Tet2^-/-^* DKO and *Tet1^-/-^, Tet2^-/-^, Tet3^-/-^*TKO lines can form “ESC-like” colonies after prolonged culture at high density in the absence of exogenously added LIF in either serum-free monolayer differentiation media, or during aggregation induced differentiation. These results may be due to an enhanced responsiveness of *Tet1^-/-^, Tet2^-/-^* DKO and *Tet1^-/-^, Tet2^-/-^, Tet3^-/-^* TKO cells to endogenously produced LIF accumulating in these high density cultures (Chambers *et al*, 2003).

Our analysis also extends prior studies by characterising redundant TET function during pluripotent state transitions and during commitment to the germline. We demonstrated that TET-deficient ESCs can differentiate normally into EpiLCs or EpiSCs, which capture *in vitro* the transitions between naïve, formative and primed pluripotent states that occur between pre- and post-implantation epiblast development (Smith, 2017; Posfai *et al*, 2021). These results offer an explanation for the *in vivo* phenotype of *Tet1^-/-^, Tet2^-/-^, Tet3^-/-^* TKO mouse embryos which are morphologically indistinguishable from wild-type until the onset of gastrulation at ∼E6.5 (Dai *et al*, 2016; Li *et al*, 2016).

TET proteins have not previously been implicated in the initial induction of PGCs from the post-implantation epiblast. While loss of TET1 is known to impair demethylation at imprinted genes and meiotic genes, this does not occur until ∼E13.5, many days after germline specification (Yamaguchi *et al*, 2012, 2013; Hackett *et al*, 2013; Hill *et al*, 2018). It was therefore unexpected that *Tet1^-/-^, Tet2^-/-^* DKO and *Tet1^-/-^, Tet2^-/-^, Tet3^-/-^* TKO cell lines should have an increased propensity to form PGCLCs during *in vitro* differentiation. Of note, TET proteins appear to be expressed in the entire epiblast of post-implantation embryos around E6.5-E7.5 (Dai *et al*, 2016; Khoueiry *et al*, 2017) and therefore do not discriminate between soma and germline. The increased PGCLC differentiation of TET-deficient cells might indicate a role for TET proteins to actively prevent germline commitment. However, it is difficult to rationalise such a role with the increasing expression of *Tet1* and *Tet2* mRNAs during the initial days of differentiation of EpiLCs to PGCLCs. Alternatively, this phenotype could be an indirect consequence of a failure of *Tet1^-/-^, Tet2^-/-^* DKO and *Tet1^-/-^, Tet2^-/-^, Tet3^-/-^* TKO cells to differentiate into somatic lineages. In this case, TET proteins could promote activation of somatic genes by demethylation of enhancers (Charlton *et al*, 2020; Ginno *et al*, 2020).

Remarkably, the enhanced germline differentiation of TET-deficient cells occurs in the absence of the otherwise requisite PGC-promoting cytokines. PGC-promoting cytokines act by rapidly downregulating *Otx2* mRNA soon after initiation of PGCLC differentiation from EpiLCs (Zhang *et al*, 2018a; Zhang & Chambers, 2019). Consequently, *Otx2*-null cells also differentiate to PGCLCs without cytokines (Zhang *et al*, 2018a). EpiLCs can also differentiate to PGCLCs without cytokines upon induced expression of NANOG (Murakami *et al*, 2016; Zhang *et al*, 2018b). As NANOG can also repress *Otx2* mRNA expression (Vojtek *et al*, 2022), the TET TKO PGCLC phenotype could occur if *Otx2* mRNA was not upregulated during the ESC-EpiLC transition. However, our analysis shows that TET-deficient EpiLCs have upregulated Otx2 mRNA. Alternatively, the similarity between the phenotypes of the *Tet1^-/-^, Tet2^-/-^* DKO, *Tet1^-/-^, Tet2^-/-^, Tet3^-/-^* TKO and *Otx2*^-/-^ lines may result from the OTX2-mediated targeting of TET proteins to somatic regulatory elements that subsequently become demethylated and active. In this scenario, ectopic OTX2 expression would not be able to rescue the TET mutant somatic block. This will be an interesting experiment for the future.

Since the lack of DNA demethylating enzymes (TETs) reported here causes a block in somatic differentiation associated with a dramatic increase in PGCLC differentiation, one might expect a lack of methylating enzymes (DNMTs) to have the opposite effect and to fully block PGCLC differentiation. However, this simple relationship does not hold. Complete loss of DNA methylation (*Dnmt1*^-/-^, *Dnmt3a*^-/-^, *Dnmt3b*^-/-^) in ESCs also causes severe defects in somatic differentiation (Tsumura *et al*, 2006; Sakaue *et al*, 2010; Schmidt *et al*, 2012), but has distinct molecular phenotypes compared to TET-deficient cells (Parry *et al*, 2023). Additionally, a recent study showed that loss of DNA methylation does not increase the number of differentiating PGCLCs but extends the period of germline competence beyond 48 hours of EpiLC differentiation (Schulz *et al*, 2024). DNMT TKO EpiLCs can also differentiate into PGCLCs in the absence of cytokines, but at an efficiency of only ∼5%, far less than observed here for *Tet1^-/-^, Tet2^-/-^, Tet3^-/-^* TKO cells. Together these results support the idea that DNA methylation (and methylation modifications) play cryptic roles in induction of correct cell fates in somatic and germline differentiation. In particular, the differences in phenotype between cells that are unable to methylate DNA and cells that cannot oxidise methylated DNA suggests quite distinct regulatory roles for intermediates in the methylation cycle in dictating choice between soma and germline, potentially by providing substrates for epigenetic “readers” (Spruijt *et al*, 2013; Iurlaro *et al*, 2013; Song *et al*, 2021; Parry *et al*, 2023). Alternatively, biological functions of TET proteins that are independent of their enzymatic activity and that occur via recruitment of the co-repressor Sin3 (Zhu *et al*, 2018; Chrysanthou *et al*, 2022; Stolz *et al*, 2022; Flores *et al*, 2023) or interplay with the Polycomb machinery (van der Veer *et al*, 2023; Chrysanthou *et al*, 2022; Huang *et al*, 2022) may be important in determining cell fate choice. Deciphering the exact mechanism by which TET proteins control gene expression at the juncture between the soma and the germline will be an important area for future investigation.

## Materials and Methods

### Cell culture

All the cell lines in this study were derived from E14Tg2a (Hooper *et al*, 1987) and incubated in a 37°C, 7% CO_2_ incubator. ESCs were routinely cultured in serum/LIF medium on gelatin coated plates. Composition of the serum/LIF medium: Glasgow minimum essential medium (GMEM; Sigma, cat. G5154), 10% fetal bovine serum, 1× L-glutamine (Invitrogen, cat. 25030-024), 1x pyruvate solution (Invitrogen, cat. 11360-039), 1× MEM non-essential amino acids (Invitrogen, cat. 11140-035), 0.1mM 2-Mercaptoethanol (Gibco ref. 31350010), 100 U/ml LIF (made in-house).

Monolayer neuronal differentiation was performed as described in (Pollard *et al*, 2006; Ying *et al*, 2003). ESCs, previously grown in serum/LIF, were dissociated using Accutase (StemPro, cat. A1110501), washed and resuspended in N2B27 medium. 100,000 cells were plated into a well a 6-well plate coated with gelatin and containing N2B27 medium. The medium was changed every 2 days until analysis. Composition of the N2B27 medium: 50ml DMEM:F12 (GIBCO cat. 12634010), 50ml Neurobasal (GIBCO cat. 21103049), 1ml 100x glutamine (Invitrogen cat. 25030024), 1ml 100x MEM non-essential amino acids (Invitrogen, cat. 11140-036), 100μl 0.1M 2-mercaptoethanol, 1ml 100x N2 supplement (GIBCO cat. 17502048), 2ml 50x B27 supplement (GIBCO cat. 17504044).

Embryoid bodies (EBs) were spontaneously formed by plating 5×10^6^ ESCs in non-adherent 5 cm dishes with GMEM medium containing 10% fetal bovine serum for 6 days. EBs were differentiated by plating on gelatin-coated wells of a 24-well plate (2 to 8 EBs per well) in GMEM medium without serum. Differentiated EBs were checked by brightfield imaging every day. In particular, the appearance of spontaneously beating clusters of cells (indicating cardiomyocyte differentiation) was noted for each well.

EpiSC lines were derived by *in vitro* differentiation from ESCs (Guo *et al*, 2009). 3×10^4^ ESCs were plated in a well of a 6-well plate with serum/LIF medium (see composition above). After 24h, the medium was switched to N2B27 medium (see composition above) supplemented with 20ng/ml Human Activin A (PeproTech, cat. 120-14E) and 10ng/ml Human Fgf basic (R&D Systems, cat. 233-FB-025/CF). Cells were submitted to daily media changes and passaged at day 5 of the protocol in 6-well plates coated with 7.5μg/mL bovine fibronectin. Cells were maintained in N2B27 medium supplemented with Activin/Fgf and passaged every 2-3 days. Homogenous EpiSCs were derived within 10 passages.

For 2i/LIF ESC culture, N2B27 medium was supplemented with 0.4μM PD0325901 (Axon, cat. 1408), 3μM CHIR99021 (Axon, cat. 1386) and 1,000 U/ml ESGRO LIF (Millipore, cat. ESG1106). EpiLC differentiation and PGCLC induction were performed as previously described (Hayashi *et al*, 2011; Hayashi & Saitou, 2013; Zhang *et al*, 2018a). ESCs were adapted to 2i/LIF culture for at least 3 passages on poly-L-ornithine (Sigma, cat. P3655) and laminin (BD Biosciences, cat. 354232) coated wells of a 6-well plate. To obtain EpiLC, 1.0×10^5^ ESCs were plated on a well of 12-well plate pre-treated with 16.6µl/ml fibronectin (Millipore, cat. FC010) and containing EpiLC medium: N2B27 medium (see composition above) supplemented with 20ng/ml Human Activin A (PeproTech, cat. 120-14E), 12ng/ml Human Fgf basic (R&D Systems, cat. 233-FB-025/CF) and 1% knock-out serum replacement (KOSR; Gibco, cat. 10828-028). After 2 days, EpiLCs were collected for analysis or submitted to PGCLC differentiation.

To obtain PGCLCs, 1.5×10^5^ EpiLCs were resuspended in 5ml (3×10^4^ cells/ml) of GK15 medium (GMEM supplemented with 15% KOSR) supplemented with 50 ng/mL bone morphogenetic protein (BMP) 4, 50 ng/mL BMP8a, 10 ng/mL stem cell factor (SCF), 10 ng/mL epidermal growth factor (EGF), and 1,000 U/mL LIF added before replating 100µl per well of a U-bottom 96-well plate (Thermo Fisher Scientific, cat. 268200). For cytokine-free differentiation, EpiLCs were similarly dissociated and resuspended in 5ml of GK15 medium (3×10^4^ cells/ml) without aforementioned cytokines. Both cells with and without cytokines were cultured for 6 days before flow cytometry analysis.

### Flow cytometry

Flow cytometry analysis was performed as previously described (Zhang *et al*, 2018a, 2018b). Cells were dissociated using trypsin and neutralized in PBS/10% serum. Cell pellets were collected by centrifugation and resuspended in 100µl PBS/2% serum supplemented with Alexa Fluor 647 anti-CD15 (SSEA-1) (Biolegend, cat. 125608) and Phycoerythrin (PE) anti-CD61 (Biolegend, cat. 104307) antibodies diluted 1/200 and 1/600, respectively. Following a 20min incubation at 4°C, cells were washed twice in 1ml PBS/10% serum before flow cytometry analysis on a BD LSR Fortessa instrument. Cells were first gated based on morphology (FSC/SSC), followed by selection for singlets based on linear correlations between FSC-area and FSC-height. Live cells were then gated based on exclusion of DAPI to indicate cell membrane integrity. Gates for SSEA-1 and CD61 were determined using unstained cells and cells stained with a single antibody.

### Self-renewal assays

ESCs, previously grown in serum/LIF, were collected by trypsinisation, resuspended in GMEM medium and counted. 600 cells were plated in gelatin-coated wells of 6-well plate containing complete GMEM medium with 10% fetal bovine serum (see composition above) with or without LIF. Following 7 days of culture, cells were washed in PBS and incubated for 1 minute in fixative solution made by mixing 25ml of citrate solution (18mM citric acid, 9mM sodium citrate, 12mM NaCl), 8ml of formaldehyde solution (37% v/v in water) and 65 ml of acetone. Fixed cells were washed in distilled water and stained for alkaline phosphatase (AP) expression using a leukocyte alkaline phosphatase kit (Sigma, cat. 86R-1KT). Colonies were counted and categorised according to their morphology and alkaline phosphatase staining.

### Knockout of *Tet1/2/3* by CRISPR/Cas9

To knockout endogenous *Tet1/2/3* alleles, two double-strand breaks were simultaneously generated using Cas9 and two synthetic guide RNAs targeting the start and stop codon of the open reading frame of the *Tet* family gene of interest, respectively. Importantly, we checked that the excised regions contained only *Tet* family genes and no other open reading frames or functional elements annotated in the genome databases (Ensembl, UCSC Genome Browser). Guide RNAs were designed (http://crispr.mit.edu/) and cloned into Cas9/gRNA co-expression plasmids carrying a fluorescent reporter (Addgene PX458 containing EGFP, or a custom-made version containing mCherry).

1×10^6^ ESCs were co-transfected with two Cas9/gRNA plasmids targeting the start and stop codon of *Tet* open reading frame, respectively, using Lipofectamine 3000 (Thermo Fisher Scientific, cat. L3000008) and following manufacturer’s instructions. After 48h, single ESCs expressing high levels of Cas9/gRNAs were selected by FACS sorting (using the EGFP/mCherry fluorescent reporter) and plated into wells of a 96-well plate coated with gelatin and containing serum/LIF medium supplemented with antibiotics (Penicillin-Streptomycin) to avoid contamination. ESC clones were expanded in 24-wells plates and genomic DNA was extracted (Qiagen, cat. 69506) for genotyping *Tet* alleles. PCR genotyping was performed using primers pairs specifically amplifying wild-type or knockout *Tet* alelles. Correctly targeted ESC clones were expanded, and frozen aliquots were transferred to liquid nitrogen tanks for long term storage.

### List of gRNAs used for generating *Tet* knockout alleles

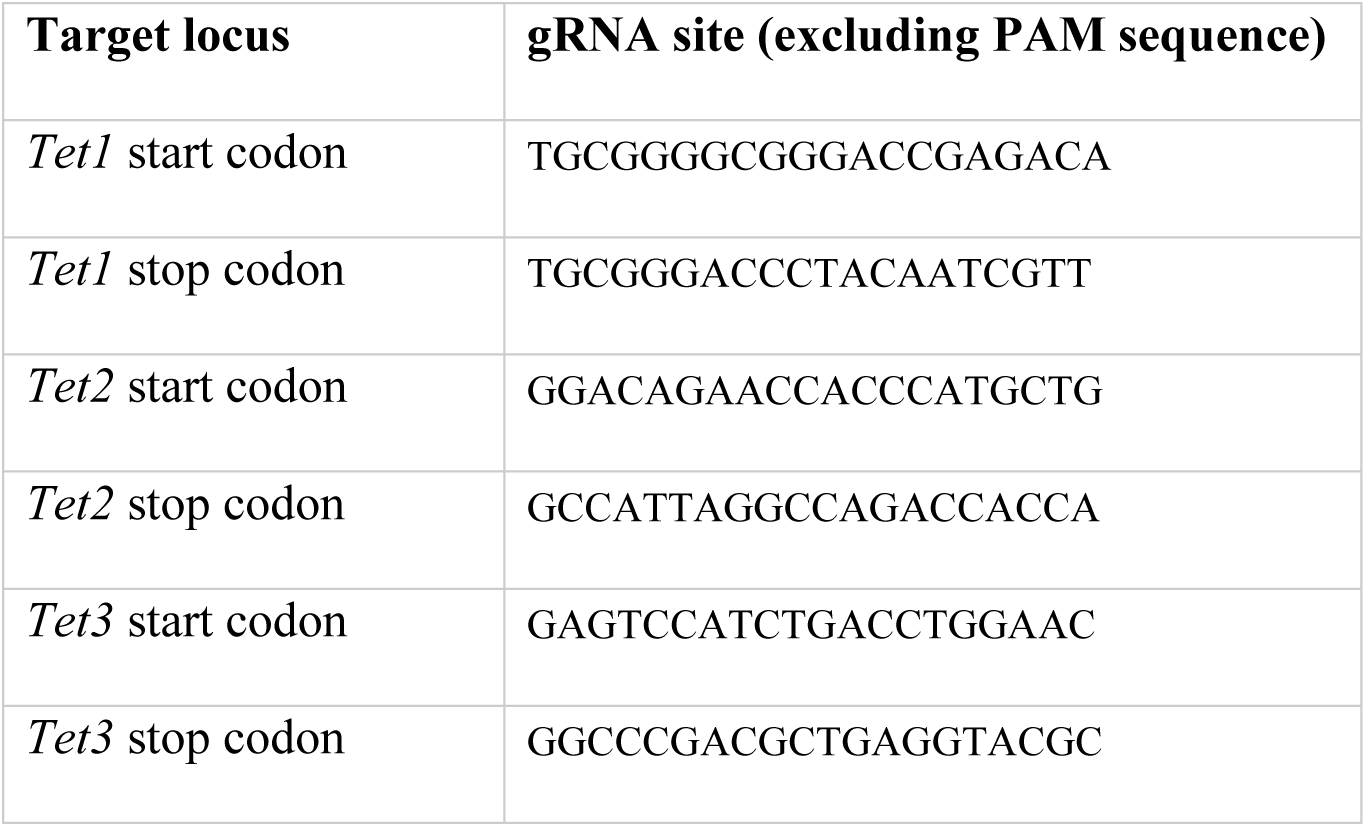

### List of primers used for genotyping *Tet* knockout alleles

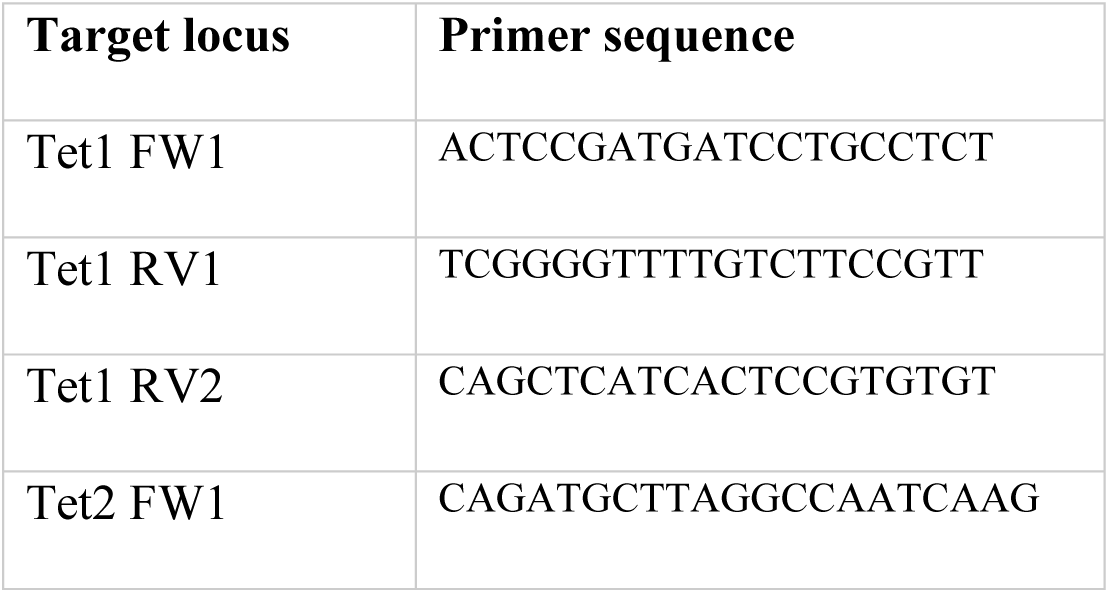
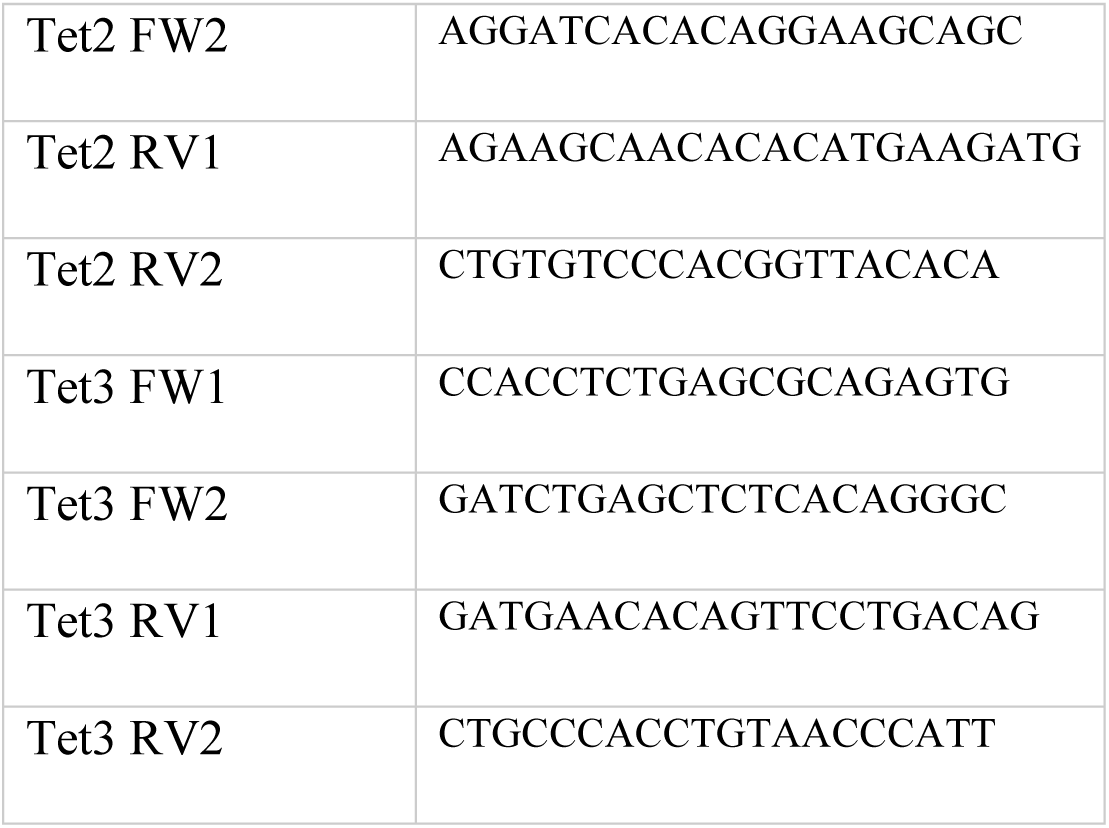

### Immunofluorescence analysis and microscopy

For brightfield analyses, live cells were imaged using an inverted microscope (MOTIC AE2000) and the software Volocity.

For immunofluorescence analyses, cells were washed with PBS and fixed with 4% PFA for 10 min at room temperature. After fixation, cells were washed with PBS and permeabilised with a solution of PBS containing 0.3% (v/v) TritonX100 for 10min at room temperature. Samples were blocked in blocking buffer (PBS supplemented with 0.1% (v/v) TritonX100, 1% (w/v) BSA and 3% (v/v) serum of the same species as the secondary antibodies were raised in) for 1h at room temperature. Following blocking, samples were incubated with primary antibodies diluted in blocking buffer overnight at 4°C. After 4 washes with PBS containing 0.1% (v/v) Triton X-100, samples were incubated with fluorescently-labelled secondary antibodies diluted in blocking buffer for 1hr at room temperature in the dark. Cells were washed 4 times with PBS containing 0.1% (v/v) TritonX100. DNA was stained with 4’,6-diamidino-2-phenylindole (DAPI) for 5 min at room temperature. Cells were washed with PBS for 5 min. Samples were imaged by fluorescence microscopy (Nikon Ti-E). Images were analysed and processed using the software Fiji. To allow comparative analyses between cell lines, samples were imaged, processed and analysed in parallel.

### List of antibodies used in this study

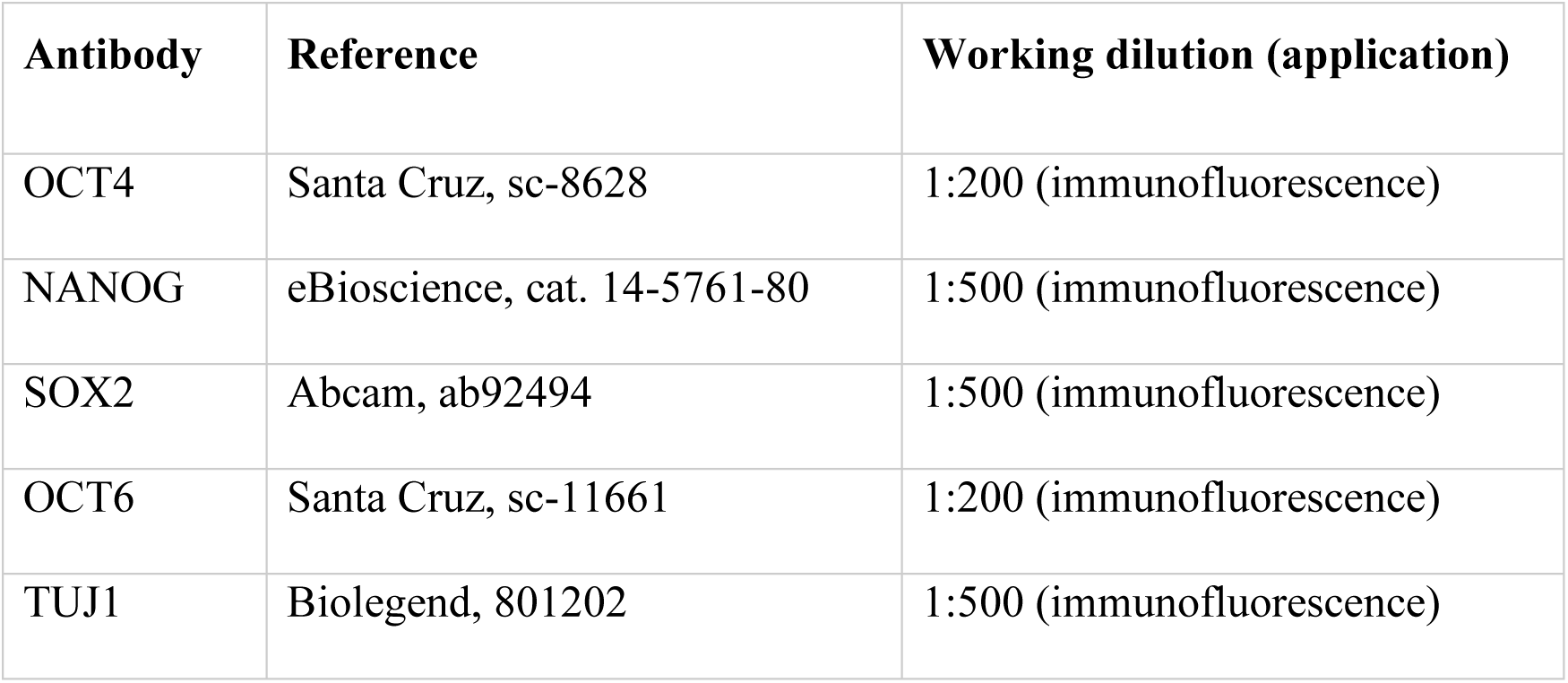

### RT-qPCR analysis

Total RNA was isolated using the RNeasy Plus Mini kit (Qiagen, cat. 74136), following manufacturer’s instructions. The quantity and purity of RNA samples were determined using a micro-volume spectrophotometer (Nanodrop, ND-1000). RNA was reverse-transcribed with SuperScript III (Invitrogen, cat. 18080044) using random hexamer oligonucleotides, following manufacturer’s instructions. Triplicate qPCR reactions were set up with the Takyon SYBR MasterMix (Eurogentec, cat. UF-NSMT-B0701) and analysed using the Roche LightCycler 480 machine. For all qPCR primer pairs, standard curves were performed to assess the amplification efficiency and melting curves were generated to verify the production of single DNA species.

### List of primers used for RT-qPCR

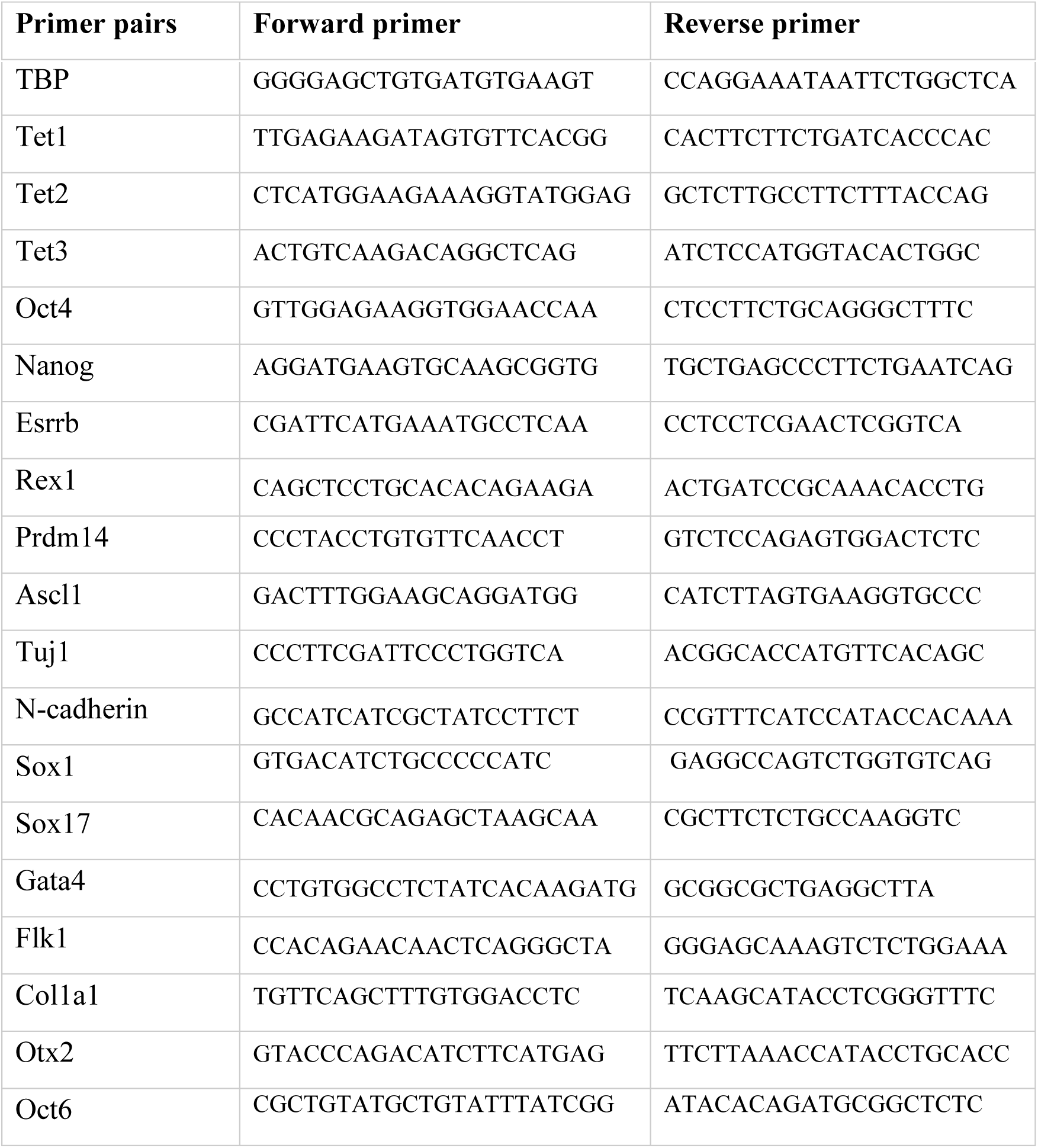

### RNA-seq analysis

RNA extraction was performed using the Direct-zol RNA MiniPrep kit (Zymo Research, cat. R2052) following the manufacturer’s instructions. RNA quality was assessed by using Tapestation 2200 (Agilent) and the ratio of RNA and DNA concentrations were determined using QUBIT The NEBNext rRNA Depletion kit (Human/Mouse/Rat) (cat. E6310) was used with 100ng of total RNA. rRNA-depleted RNA was then DNase treated and purified using Agencourt RNAClean XP beads (Beckman Coulter Inc, cat. 66514) and the NEBNext Ultra II Directional RNA library prep kit for Illumina (cat. E7760) was used to generate final libraries.

Sequencing was then performed on the NextSeq 2000 platform (Illumina Inc, cat. 20038897). FASTQ files were generated in Illumina’s BaseSpace and quality trimming was performed with TrimGalore 0.5.0 and cutadapt 1.16 (Martin, 2011). Reads were aligned to the mouse genome (mm10) using STAR 2.7.1a (Dobin *et al*, 2013). Count matrices were then generated using subread 1.5.2 and differential expression analysis was performed by Deseq2 (Love *et al*, 2014).

## Data availability

Original RNA-seq data are deposited at the NCBI Gene Expression Omnibus (GEO) with accession number GSE273732.

## Acknowledgements

We thank Bertrand Vernay for microscopy support, Claire Cryer and Fiona Rossi for assistance with flow cytometry, the Genetic Core Sequencing Service (WGH, Edinburgh) for library preparation and sequencing of RNA-seq samples, Andrea Corsinotti for qPCR primers and Val Wilson and Steve Pollard for comments on the manuscript. This work was funded by UK Medical Research Council Grants MR/L018497/1 and MR/T003162/1 to IC. EB was supported by a Marie Sklodowska-Curie fellowship (H2020-MSCAIF-2018/843879). RP was supported by a MRC PhD studentship, SBG by an MRC Precision Medicine PhD studentship and TT by a Darwin Trust of Edinburgh PhD studentship.

## Author contributions

RP and IC conceived the project. RP designed, performed and analysed somatic differentiation experiments with contributions from ET (immunofluorescence), TT (EpiLC differentiation) and DC (PGCLC and EB differentiation). EB, SGB and MZ performed and analysed PGCLC differentiation. SGB analysed RNA-seq. RP, EB, SGB and IC wrote the manuscript with editing by all authors.

## Supplementary figure legends

**Expanded View Figure 1.**
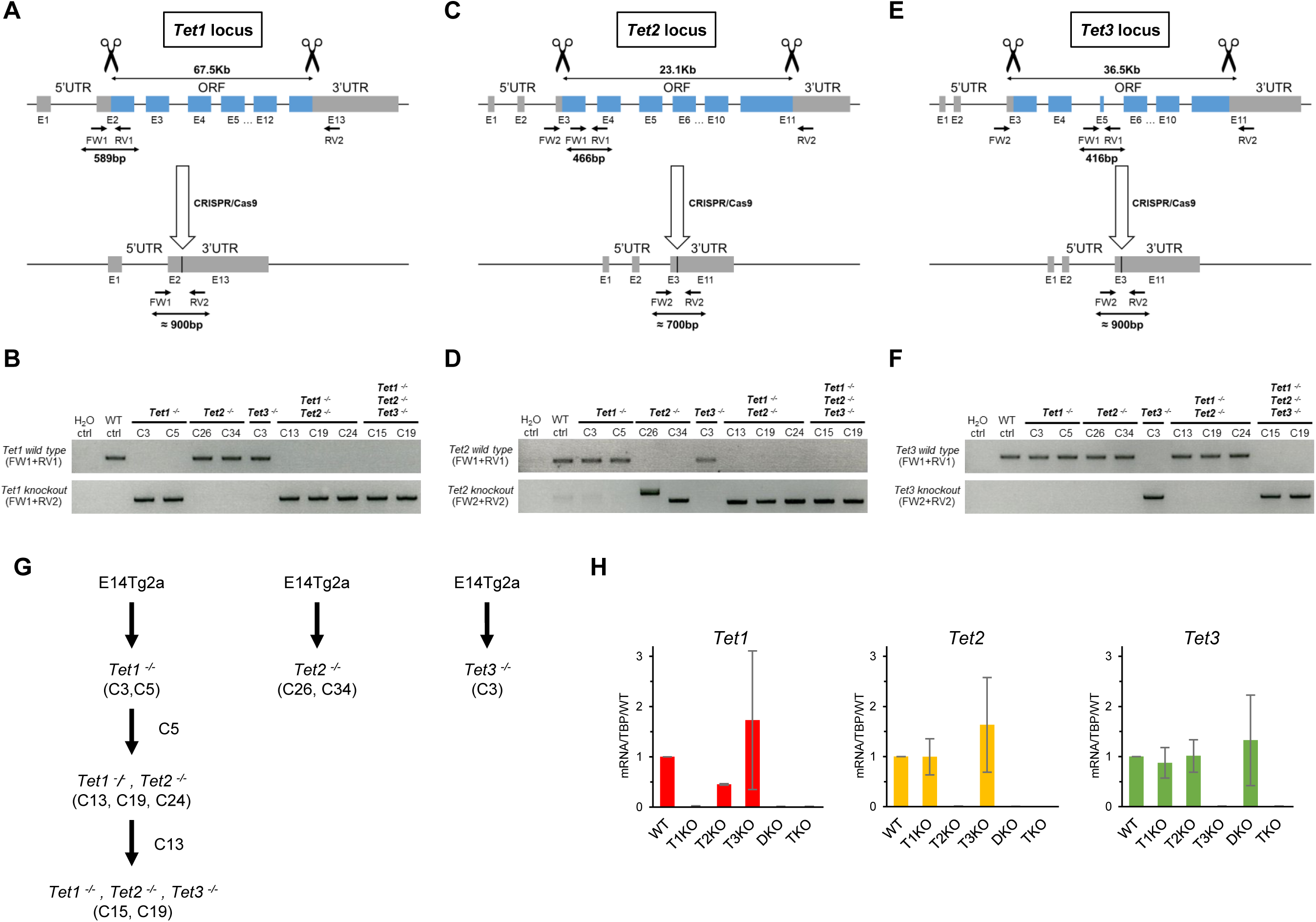
Genetic deletion of *Tet1/2/3* open reading frames by CRISPR/Cas9. For all loci the general targeting strategy is indicated (A, C, E) with two gRNAs (scissors) designed close to the start and stop codon, respectively. The position of genotyping primers and the sizes of PCR products are indicated. Open reading frames are indicated in blue and untranslated regions (UTRs) in grey. (B, D, F) agarose gel analysis of PCR genotyping of *Tet1, 2 and 3* alleles in all cell lines, with the PCR reaction indicated on the left, and the deduced genotype at the top. **A.** To knockout *Tet1*, wild-type E14Tg2a ESCs were co-transfected with Cas9 and two gRNAs targeting the *Tet1* start and stop codons, respectively. **B.** Two *Tet1^-/-^* clones carrying deletions of both *Tet1* alleles (C3 and C5) were obtained, as demonstrated by the presence of a PCR product for the knockout allele (FW1+RV2) and the absence of a PCR product for the wild-type allele (FW1+RV1). **C.** *Tet2* knockout and *Tet1, Tet2* double knockout (hereafter referred to as DKO) cell lines were generated from E14Tg2a and *Tet1^-/-^* C5 ESCs using the strategy shown. **D.** Clones (C26, C34) lacking *Tet2* alleles and clones lacking both *Tet1* and *Tet2* alleles (C13, C19, C24) were generated. **E.** *Tet3* knockout and *Tet1, Tet2, Tet*3 triple knockout (hereafter referred to as TKO) ESCs lacking all six *Tet* alleles were generated from E14Tg2a and *Tet1^-/-^*, *Tet2^-/-^* C13 ESCs as illustrated. **F.** One clone lacking *Tet3* alleles (C3) and two TKO clones lacking all alleles for *Tet1, Tet2* and *Tet3* (C15, C19) were obtained. WT ctrl. Wild-type E14Tg2a ESCs (parental cell line), H_2_O ctrl. Control sample with no DNA template. Targeting events leading to the generation of single and combined *Tet* knockout ESC lines are detailed in Figure S1. **G.** Summary of the *Tet* knockout ESC lines generated in this study and their interrelationships (see Figure 1). Homozygous clone numbers are indicated within brackets. New rounds of targeting are indicated with arrows. **H.** *Tet1*, *Tet2* and *Tet3* mRNA levels in the indicated ESC lines. mRNA levels were quantified by RT-qPCR, normalised to TBP mRNA levels, and expressed relative to levels in wild-type E14Tg2a ESCs (mRNA/TBP/WT ESC). Error bars: standard deviation in 2 independent replicate experiments.

**Expanded View Figure 2.**
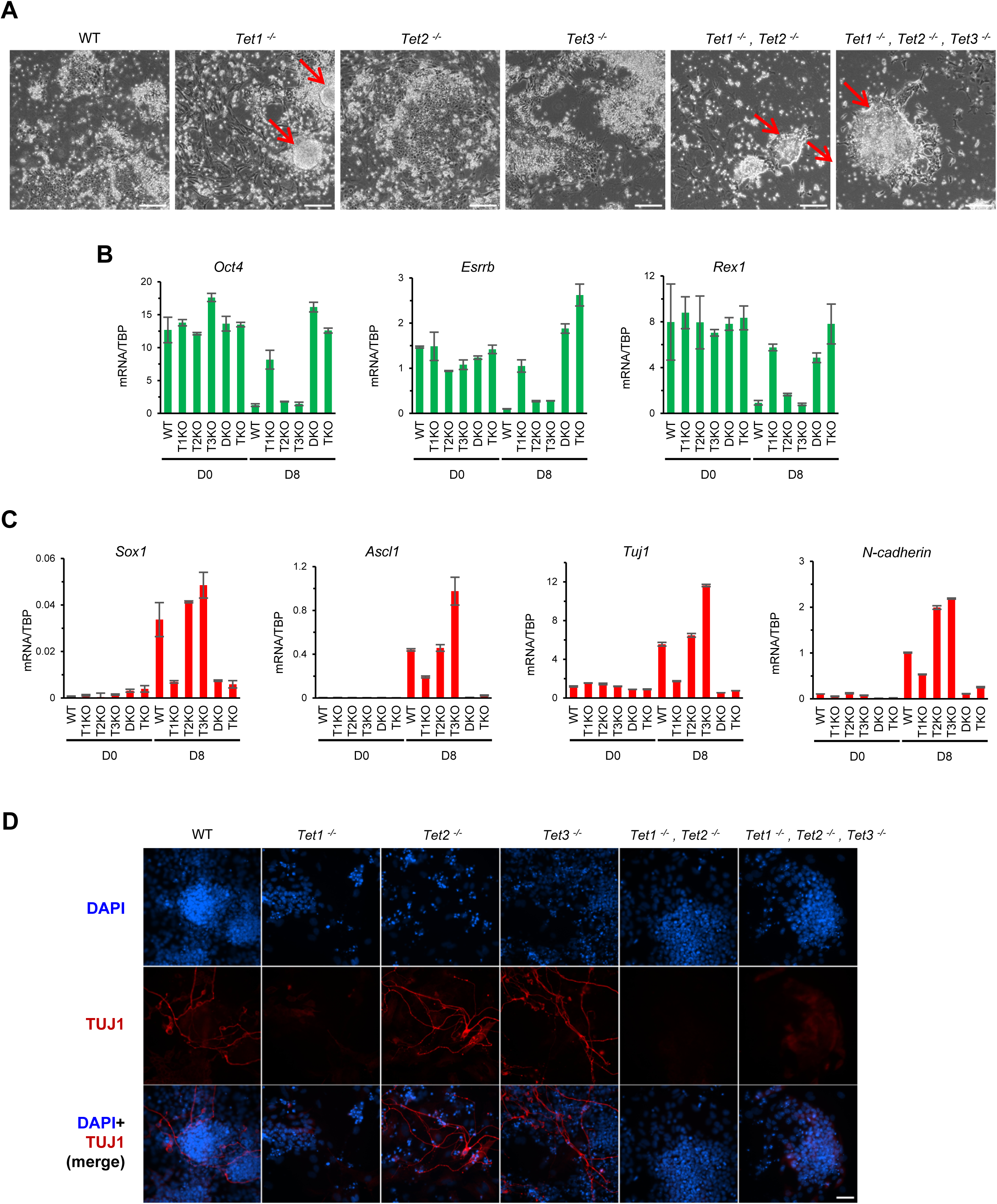
Phenotype of TET-deficient cells during monolayer neural differentiation. **A.** Phase contrast images of the indicated live cell lines after monolayer neural differentiation for 6 days. Colonies retaining an undifferentiated morphology are indicated by red arrows. Scale bars: 100µm. **B.** mRNA levels of pluripotency factors in the indicated cell lines cultured as ESCs (D0) in serum/LIF or after 8 days of monolayer neural differentiation (D8). mRNA levels were quantified by RT-qPCR and normalised to TBP mRNA levels. Data are from a representative of 2 independent experiments (error bars: standard deviation in 3 technical replicates). **C.** mRNA levels of neural markers in the indicated cell lines cultured as ESCs (D0) in serum/LIF or after 8 days of monolayer neural differentiation (D8). mRNA levels were quantified by RT-qPCR and normalised to TBP mRNA levels. Error bars: standard deviation in 3 technical replicates. **D.** Immunofluorescence for Tuj1 in the indicated cell lines cultured in N2B27 medium for 8 days. Scale bar: 50µm.

**Expanded View Figure 3.**
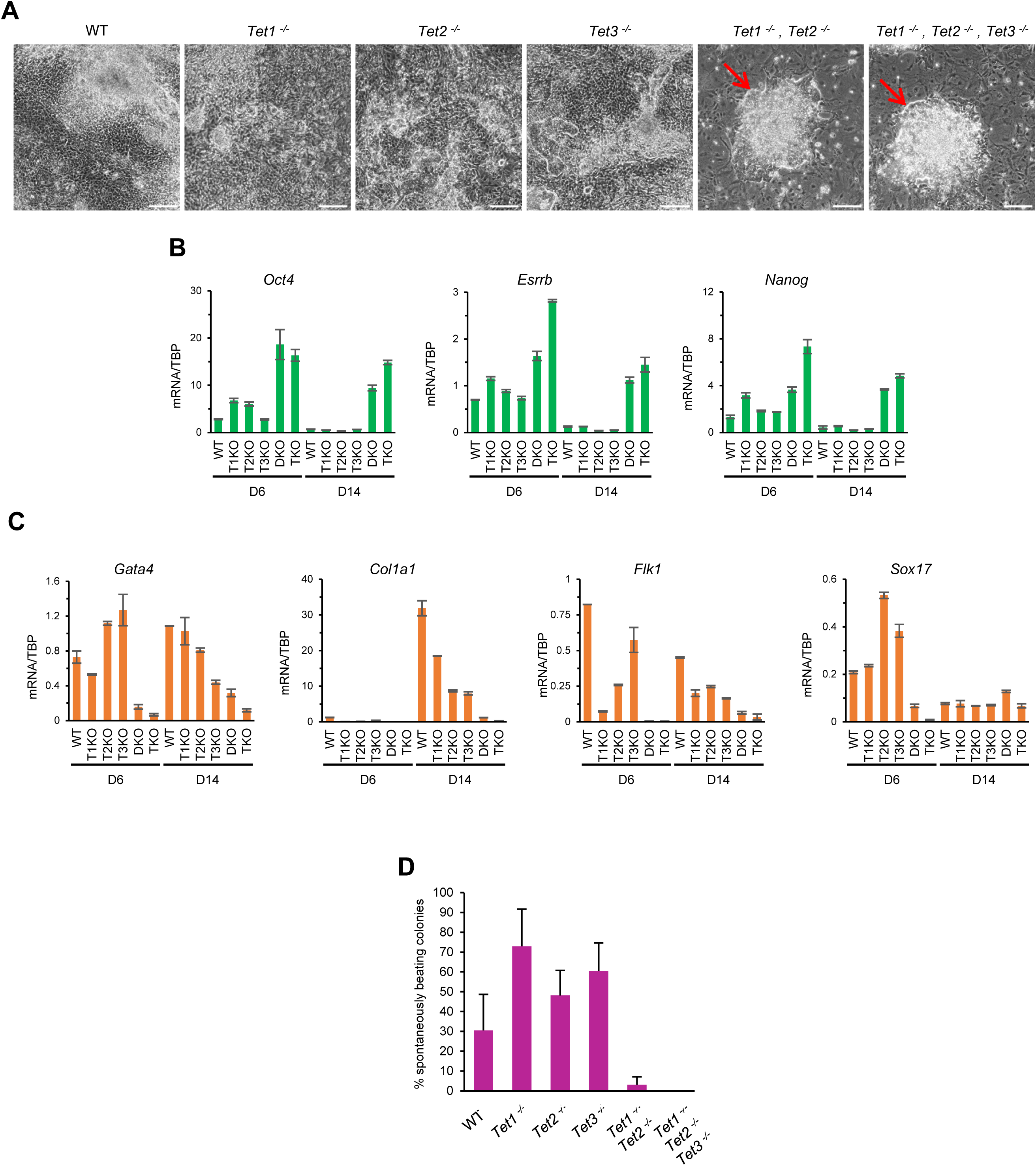
Phenotype of TET-deficient cells during embryoid body differentiation. **A.** Brightfield images of the indicated live cell lines following 13 days of embryoid bodies (EB) differentiation. Colonies retaining an undifferentiated morphology are indicated by red arrows. Scale bars: 100µm. **B.** mRNA levels of pluripotency factors in the indicated cell lines following embryoid body formation (D6) and after a further 8 days of differentiation as outgrowths (D14). mRNA levels were quantified by RT-qPCR and normalised to TBP mRNA levels. Data are from a representative of 2 independent experiments (error bars: standard deviation in 3 technical replicates). **C.** mRNA levels of lineage markers in the indicated cell lines following embryoid body formation (D6) and after a further 8 days of differentiation as outgrowths (D14). mRNA levels were quantified by RT-qPCR and normalised to TBP mRNA levels. Data are from a representative of 2 independent experiments (error bars: standard deviation in 3 technical replicates). **D.** Proportion (%) of spontaneously beating colonies following 13 days of EB differentiation with the indicated cell lines (n=4).

**Expanded View Figure 4.**
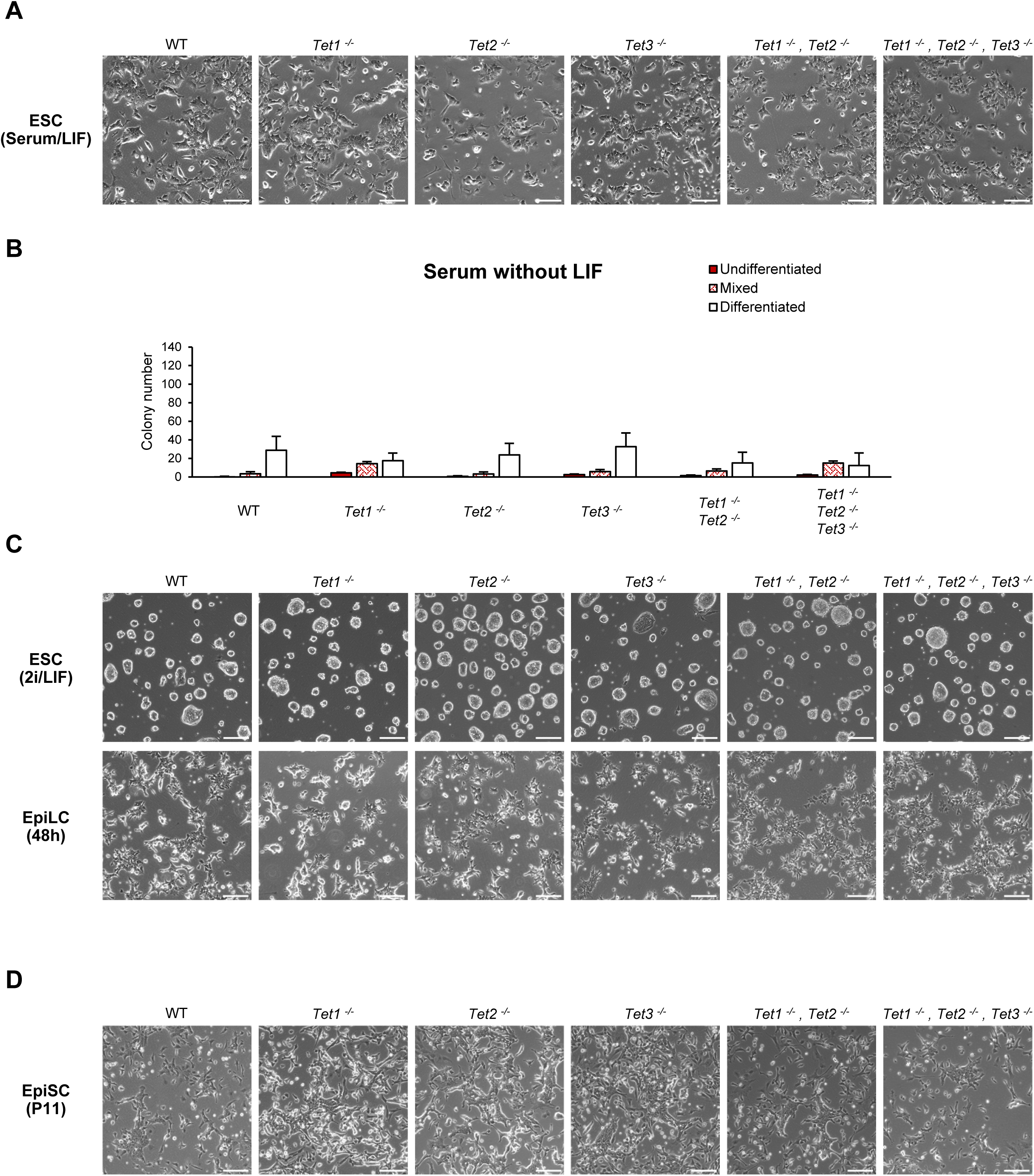
Phenotype of TET-deficient in different pluripotent states. **A.** Phase contrast images of the indicated live ESC lines cultured in serum/LIF. Scale bars: 100µm. **B.** Alkaline phosphatase staining of the indicated ESC lines following plating at clonal density in serum-containing medium without LIF for 7 days. Colonies were counted and categorised according to their alkaline phosphatase staining. Error bars: standard deviation in 4 independent replicate experiments. **C.** Brightfield images of the indicated live ESC lines cultured in 2i/LIF and following 48h of EpiLC differentiation in activin/FGF. Scale bars: 100µm. **D.** Phase contrast images of the indicated live EpiSC lines (passage 11) cultured in activin/FGF. Scale bars: 100µm.

**Expanded View Figure 5.**
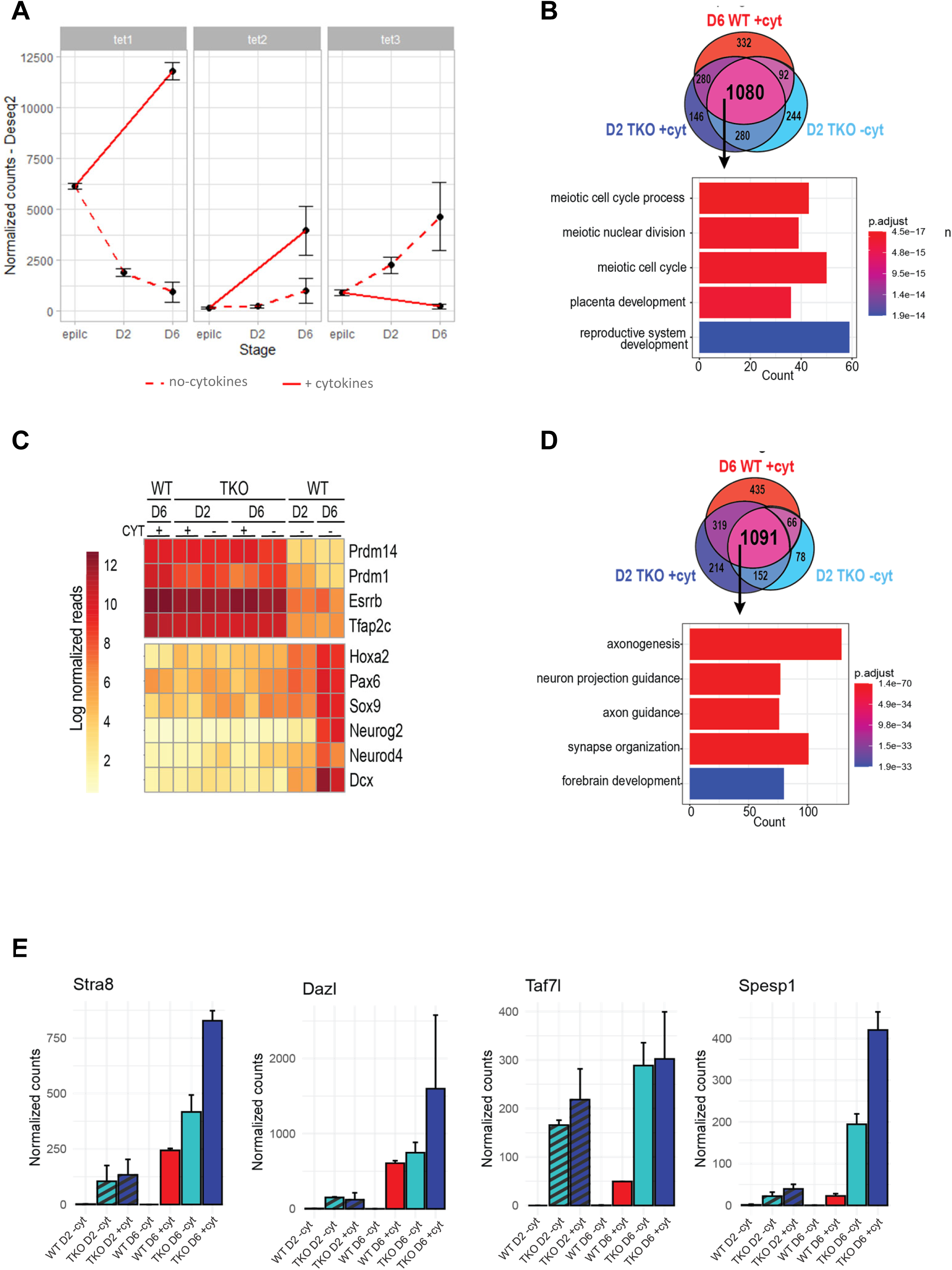
Timecourse RNA-seq analysis of TET triple knockout PGCLC differentiation. **A.** Expression dynamics of *Tet1, Tet2* and *Tet3* in wild type cells during PGCLC differentiation in the presence and absence of cytokines. Mean ±SD of RNA-seq reads normalized by DESeq2 of two biological replicates. **B.** Overlap of all genes detected as significantly upregulated in day 6 wild-type PGCLCs and day 2 *Tet1^-/-^, Tet2^-/-^, Tet3^-/-^* TKO (+/-cytokines) compared to day 6 wild-type cells differentiated without cytokines (q value < 0.05, Log2 fold change >2). Gene ontology enrichment results shown for genes shared across all three samples. **C.** Log normalized reads of germline and somatic markers. **D.** Overlap of all genes detected as significantly downregulated in day 6 wild-type PGCLCs and day 2 *Tet1^-/-^, Tet2^-/-^, Tet3^-/-^* TKO (+/-cytokines) compared to day 6 wild-type cells differentiated without cytokines (q value < 0.05, Log2 fold change < -2). Gene ontology enrichment results shown for genes shared across all three samples. **E**. Expression of late germline markers. Mean ±SD of RNA-seq reads normalized by DESeq2 of two biological replicates.

